# Met-signaling Controls Dendritic Cell Migration by Regulating Podosome Formation and Function

**DOI:** 10.1101/2021.04.28.441791

**Authors:** Ahmed E.I. Hamouda, Carmen Schalla, Antonio Sechi, Martin Zenke, Thomas Hieronymus

## Abstract

Signaling by the HGF receptor/Met in skin-resident Langerhans cells (LC) and dermal dendritic cells (dDC) is essential for their emigration toward draining lymph nodes upon inflammation-induced activation. Here we addressed the role of Met-signaling in distinct steps of LC/dDC emigration from skin by employing a conditional Met-deficient mouse model (Metf^lox/flox^). We found that Met deficiency severely impaired podosome formation in DC and concomitantly decreased proteolytic degradation of gelatin. Accordingly, Met-deficient LC failed to efficiently cross the extracellular matrix (ECM) rich basement membrane between epidermis and dermis. We further observed that Met-signaling by HGF reduced adhesion of bone marrow-derived LC to various ECM factors and enhanced the motility of Met-signaling competent DC in a 3D collagen matrix, which was not the case for Met-deficient LC/DC. We found no impact of Met-signaling on the integrin-independent amoeboid migration of DC in response to the CCR7 chemokine CCL19. Collectively, our data show that the Met-signaling pathway regulates the migratory properties of DC in HGF-dependent and HGF-independent manners.

## Introduction

The development and immunological functions of dendritic cells (DCs) are intricately coupled to their motile and migratory properties (Bancherau and Steinman, 1998; Alvarez et al. 2008). DC precursors develop at sites of hematopoiesis throughout embryogenesis and adult life. Then they immigrate into peripheral and lymphoid tissues where they differentiate into immature DCs to act as sentinels of the immune surveillance system (Merad et al., 2013). The motile properties of DC allow them to sample the environment for antigens (Ags). Following Ag capture, DCs are activated and migrate via lymphatics to lymphoid organs where they present processed Ags to naïve T cells to establish an adaptive immune response (Bancherau and Steinman, 1998; Merad et al., 2013). Thus, throughout a DC life cycle, migration and homing are tightly regulated by the specific expression of chemokines and chemokine receptors, adhesion molecules, and homing receptors, among others, thereby warranting the selective placement of DC at specific functional sites.

In skin, different DC subsets constitute the first immune barrier for invading pathogens and contact allergens, namely Langerhans cells (LC), the only DC subset of the epidermis and various dermal DC substes (dDC) that differ in development and function (Henri, et al., 2010; Merad et al., 2013, Kashem et al., 2017). Emigration of skin-resident LC/dDC towards the skin draining lymph nodes (sdLN) upon activation requires a multitude of tissue remodeling capacities, including expression regulation of adhesion molecules, which allow (i) their detachment from surrounding tissues, (ii) adherence to and migration through the extracellular matrix (ECM) of the interstitial space, and (iii) crossing tissue boundaries. Proinflammatory cytokines, such as TNFα, have been recognized as important mediators of DC mobilization, leading to the emigration from peripheral tissues (Alvarez et al., 2008; Merad et al., 2013).

Initial studies by our group identified hepatocyte growth factor (HGF) as a potent elicitor of LC emigration from mouse epidermis like TNFα (Kurz et al., 2002). We also found the receptor tyrosine kinase (RTK) Met, the cognate high-affinity transmembrane receptor of HGF, to be expressed on skin DC populations, including LC and dDC, and on bone marrow (BM)-derived DC (BMDC) (Kurz et al., 2002, Baek et al., 2012). Met was originally identified as an oncogene that can induce invasive growth and metastasis of tumor cells (Birchmeier et al., 2003; Trusolino et al., 2010). Met-signaling is indispensable during embryonic development and is critically involved in wound healing. Moreover, Met has also been implicated in hematopoiesis and in immune regulation, as expression of Met was found in hematopoietic progenitor cells, macrophages, neutrophils, B cells, DCs, and subsets of T-cells (Molnarfi et al., 2015; Ilangumaran et al., 2016; Sagi and Hieronymus 2018). In addition, HGF was found to regulate various functions of immune cells, including cytokine production, migration, and adhesion. However, the mechanisms by which Met-signaling impact on immunological responses are only scarcely explored and therefore not yet fully understood.

Clear evidence for the role of Met-signaling in skin DCs came again from our studies using a conditional Met-knockout mouse model (Borowiak et al., 2004) in which DC migration and DC-dependent contact hypersensitivity (CHS) reactions were addressed (Baek et al. 2012). Activated Met-deficient skin-resident DC failed to emigrate toward the sdLN despite displaying an activated phenotype. Consequently, Met-deficiency resulted in strongly impaired CHS reactions in response to contact allergens. Additionally, Met-signaling was found to be essential for BMDC migration through ECM dense Matrigel, which requires matrix degradation by the proteolytic activities of matrix metalloproteinase (MMP). Indeed, proteolytic activities of both MMP-2 and MMP-9 were found to be regulated by Met-signaling in BMDC (Baek et al., 2012), which is in line with previous studies that revealed a critical role of MMP2 and MMP9 in LC/DC migration (Kobayashi et al., 1999; Ratzinger et al., 2002; Yen et al., 2008; He et al., 2018).

In this study, we aimed to investigate in which distinct steps of LC/dDC emigration from skin Met-signaling is involved. By employing a conditional Met-deficient mouse model (Metf^lox/flox^), we show that the absence of Met-signaling did not affect the downregulation of E-cadherin and epithelial cell adhesion molecule (EpCAM) expression after activation, allowing their detachment from surrounding tissue and their migration through the epidermis. However, Met-deficient LC failed to cross the basement membrane towards the dermis. We also observed dDC to be impaired in their migration through the dermis. However, we found no impact of Met-signaling on the integrin-independent amoeboid migration of DC in response to the c-c chemokine receptor 7 (CCR7) ligand CCL19. Meanwhile HGF stimulation resulted in enhanced motility of Met-signaling competent DC but not Met-deficient DC. We further demonstrate that Met-signaling affects podosome formation in DC, which is severely impaired by Met-deficiency, resulting in diminished gelatin degradation. Additionally, Met-signaling by HGF reduced adhesion of BM-derived LC to various ECM factors, namely fibronectin, laminin and collagen IV. Together, these findings show that the Met-signaling pathway regulates the migratory properties of DC in HGF-dependent and HGF-independent manners.

## Materials and Methods

### Animals

Transgenic mice expressing Cre under the control of the IFN-inducible Mx promoter (B6.Cg-Tg(Mx1-Cre)1Cgn/J), C57BL/6 mice heterozygous for the Met knockout allele (Met^+/−^) and Met^flox//flox^ mice (Borowiak et al., 2004) were maintained under specific pathogen-free conditions in the central animal facility of the Rheinisch-Westfälische Technische Hochschule University Hospital Aachen. All animal experiments were approved by local authorities in compliance with the German animal protection law and EU guidelines (2010/63/EU) for animal protection. Met^+/−^ mice were crossed to Mx-cre mice and Met^flox//flox^ mice were interbred with Mx-cre^+^ x Met^+/−^ mice to generate Mx-cre x Met^+/flox^ (Met-wild type [WT]) and Mx-cre x Met^flox/−^ progeny. Mx-cre^+^ x Met^flox/−^ offspring (5-7 wk) received five injections of 300 μg polyinosinic/polycytidylic acid (pI:pC, Sigma-Aldrich) in PBS i.p. at daily intervals for excision of the floxed exon of Met to generate Met-deficient Met^Δ/−^mice (Met-knockout [KO]) mice. Further functional studies in Met-KO and Met-WT mice were performed ≥4 wk after the last pI:pC injection.

### Flow cytometry and fluorescence activated cell sorting

Flow cytometry analysis was performed as previously described (Kurz et al., 2000) on a FACSCanto II using FACSDiva software (BD Biosciences). Fluorochrome-conjugated antibodies to stain for CD3 (clone 145-2C11), CD11b (M1/70), CD11c (N418), CD40 (1C10), CD45 (30-F11), CD80 (1G10), CD86 (GL1), MHC class II (MHC-II, I-A/I-E; clone M5/114.15.2), CD103 (2E7), EpCAM/CD326 (G8.8) and biotin-conjugated antibody for staining of CD24 (M1/69) were purchased from eBioscience. PE-labeled antibody for staining of E-cadherin (114420) was obtained from R&D Systems. APC-conjugated anti-Sirpα/CD172a (P84) was from BD Biosiences. Fluorochrome-conjugated antibodies for detection of Langerin/CD207 (4C7) and XCR1 (ZET) were purchased from BioLegend. To account for fluorescence spillover, single-stained compensation beads (BD Biosciences) were recorded for each experiment. Sorting of immunofluorescently-labeled cells was done using a FACSAria II cell sorter (BD Biosciences). Acquired data were analyzed using FlowJo software (FlowJo LLC, USA).

### LC/DC isolation from murine skin

Single cell suspensions of murine epidermis and dermis were generated as previously described (Stoitzner et al., 2010). Briefly, mouse ears were split into dorsal and ventral halves and allowed to float on 0.5% trypsin-EDTA (Gibco, USA) at 37°C for 25 and 45 min, respectively. Digestion was blocked by transferring the halves to a dish containing FCS (Gibco, USA) and the epidermis peeled off the dermis. The epidermal sheets were then shaken at 1400 rpm in complete RPMI 1640 medium supplemented with 10% FCS at 37°C for 30 min. Dermal sheets were digested with 1 mg/ml collagenase D (Gibco, USA) in complete RPMI 1640 medium/10% FCS for 45 min at 37°C in a thermomixer at 1400 rpm. Following digestion, dermal and epidermal single cell suspensions were passed through a 21-gauge needle and a cell strainer and assessed by flow cytometry. Alternatively, LC were isolated from epidermal single cell suspensions by FACS sorting.

### LC/DC cell culture

A two-step amplification and differentiation protocol to generate BM-derived DC (BMDC) using GM-CSF was followed as described previously (Hieronymus et al., 2005). Briefly, BM cells were flushed out of mouse femur and tibia. BM progenitor cells were cultured at a density of 2 × 10^6^ cells/ml in complete RPMI 1640 medium /10% FCS, 30 U/ml murine SCF (CHO KLS C6 cells), 25 ng/ml human Flt3-ligand (Peprotech, USA), 40 ng/ml recombinant human long-range IGF-1 (Sigma-Aldrich, USA), 5 ng/ml recombinant IL-6/soluble IL-6R fusion protein ((Fischer et al., 1997), kindly provided by S. Rose-John), 20 U/ml recombinant murine GM-CSF (PharmedArtis, Germany) and 10^−6^ M dexamethasone (Sigma-Aldrich, USA). Progenitor cells were amplified 5 to 11 days before differentiation into DC was induced by supplementing 200 U/ml recombinant murine GM-CSF. Culture medium was replenished every second day. Immature DC were obtained after 8 to 10 days under differentiation conditions.

LC-like cells were generated from BM as described previously (Capucha et al., 2018). Briefly, BM cells were cultured under serum-free conditions at a density of 1 × 10^6^ cells/ml in complete RPMI 1640 medium supplemented with 30 U/ml murine SCF, 50 ng/ml human Flt3-ligand, 2.5 ng/ml recombinant murine TNFα (Peprotech, USA), 200 U/ml recombinant murine GM-CSF and 10 ng/ml recombinant human TGFβ (R&D Systems, USA). Partial medium change was performed at day 3 of differentiation. Immature BMLC were obtained at day 5 of differentiation.

BMDC and BMLC maturation was induced by overnight treatment with 100 ng/ml recombinant murine TNFα (Sigma-Aldrich, USA). In some experiments, 50 ng/ml HGF (eBioscience, USA) was added during maturation.

### Ex vivo migration assay

Emigration of tissue resident cells from skin *ex vivo* (crawl out assay) was performed as previously described with minor modifications (Kurz et al., 2002; Stoitzner et al., 2010). Dorsal and ventral halves of mouse ears were separated and allowed to float split-side down on complete RPMI 1640 medium supplemented with 10% FCS, 200 U/ml GM-CSF and 100 ng/ml murine CCL19 (Peprotech, USA) at 37°C for 24-72 h.

At the end point, the ear halves were either digested and analyzed with flow cytometry or they were incubated with 20 mM EDTA for 90 min to facilitate the separation of the epidermis. Then, epidermal sheets were stained and analyzed with immunofluorescence microscopy. Cells that emigrated into the culture medium were collected, stained and analyzed by flow cytometry. Numbers of migrated cells were normalized using Dynabeads (15-μm diameter; Dynal Polymers, Norway), as previously described (Kurz et al., 2002; Beak et al., 2012)

### Immunofluorescence microscopy of epidermal sheets

Epidermal sheets were fixed in ice-cold acetone for 20 min, washed with PBS and blocked using 3% (w/v) BSA in PBS for 30 min at RT. The epidermal sheets were stained at 4°C overnight with fluorochrome conjugated antibodies against MHC-II (M5/114.15.2) and CD3 (145-2C11) diluted in blocking buffer. Following antibody staining, epidermal sheets were washed with PBS and stained with 100 ng/ml DAPI (Vector Laboratories, USA) for 5 min at RT. Finally, epidermal sheets were mounted on glass slides using DAKO fluorescence mounting medium (Agilent Technologies, USA). Images were acquired using an Axioplan 2 microscope (Zeiss, Germany) equipped with coolSNAP HQ2 camera operated with VisiVIEW imaging software (Visitron Systems, Germany). Image processing was done using Fiji (https://imagej.net/Fiji).

### Confocal microscopy

Cryosections of mouse footpad skin (25 μm thick) were fixed in ice-cold acetone on ice for 30 minutes, washed with PBS and blocked with 5% (v/v) FCS in PBS for 30 minutes at RT. The skin sections were stained with fluorochrome conjugated antibodies against MHC-II and CD49f (GoH3) at 4°C overnight. Counterstaining was done with 100 ng/ml DAPI for 5 minutes at RT. Finally, the cryosections were mounted in ProLong Gold mounting media (Molecular Probes, USA) on glass slides. Images were acquired using LSM700 confocal microscope (Zeiss, Germany) equipped with a 40 x oil immersion objective. Confocal images were processed using Fiji.

### Gelatin degradation and podosome formation assay

The gelatin degradation and podosome formation assay was performed essentially as previously described with some modifications (Díaz, 2013). Briefly, round 12 mm glass coverslips were first coated with 50 μg/ml poly-l-lysine (Sigma-Aldrich, USA) diluted in water for 20 min at RT followed by washing with PBS and fixation with 0.5 % (v/v) glutaraldehyde (Sigma-Aldrich, USA) for 15 min on ice. Next, the coverslips were coated with 0.2 mg/ml Oregon green 488-labeled gelatin (Invitrogen, USA) diluted in 2% (w/v) sucrose in PBS for 30 min in the dark at RT. After that, coverslips were incubated with 0.3 M glycine (pH 7.2) for 20 minutes at RT to quench glutaraldehyde-induced autofluorescence. The coated coverslips were washed with PBS under sterile conditions and were equilibrated by incubating them with complete RPMI 1640 supplemented with 10% FCS for 30 min at 37°C before cell seeding.

1 × 10^5^ BMDC were seeded on each coverslip in the presence of TNFα and incubated overnight at 37°C. In some experiments, BMDC were further treated with the inhibitors PD98059, SU6656 (all from Calbiochem, USA) or SU11274 (Selleck Chemicals, USA). At the end point, BMDC are fixed with 4% (w/v) PFA in cytoskeleton buffer for 15 min at RT. Then, BMDC were blocked and permeabilized by 3% (w/v) BSA/0.1% (v/v) TritonX-100 in PBS for 30 min at RT and stained for actin with Alexa594-conjugated phalloidin (BioLegend, USA) diluted in 0.3% (w/v) BSA/0.1% (v/v) TritonX-100 in PBS for 1 h at RT. After excess washing with PBS, BMDC were stained with 100 ng/ml DAPI (Vector Laboratories, USA) for 5 min at RT. Finally, the coverslips were mounted using DAKO fluorescence mounting medium (Agilent Technologies, USA). Image acquisition and processing was done using an Axioplan 2 microscope and Fiji software.

### Chemotaxis and motility assay in 3D collagen gels

3D chemotaxis and motility assays were performed as previously described with some modifications (Sixt and Lämmermann, 2011). Acid washed glass coverslips (18 × 18 mm) were attached using polydimethylsiloxane (PDMS; Sylgard 184 Silicone Elastomer Kit, Dow Chemicals, USA) to the bottom and upper side of modified tissue culture dishes that have a 12 mm round hole. This creates an approximately 0.5 mm thick chamber. Collagen G (Merck, Germany) gel is prepared according to the manufacturer’s instructions. 2 × 10^6^/ml BMDC in complete RPMI 1640 supplemented with 10% FCS are mixed 1:1 with the collagen G solution, generating a 1.6 mg/ml collagen/BMDC mixture. Two thirds of the chamber is filled with the collagen/BMDC mixture and incubated for 1 h at 37°C in an upright position, thereby preventing BMDC from settling beneath the collagen gel and promoting gelation.

For chemotaxis assay, the remaining one-third of the chamber was filled with complete RPMI 1640 supplemented with 10% FCS in the presence or absence of 0.6 μg/ml murine CCL19 (Peprotech, USA). The slowly diffusing CCL19-containing medium creates a CCL19 gradient that promotes BMDC migration towards the upper part of the chamber.

For motility assay, the remaining one-third of the chamber was filled with complete RPMI supplemented with 10% FCS in the presence or absence of 50 ng/ml HGF (eBioscience, USA) and the inhibitors PD98059 or SU11274.

Bright-field time-lapse recordings of cells within 3D collagen gels were acquired at 1 frame/min for 4 h using an inverted AxioVert 200 microscope (Zeiss, Germany) equipped with a humidified climate chamber (PeCon, Germany) adjusted at 5% CO2, 37°C and a Cascade 512B EMCCD camera (Photometrics, USA) operated with IPlab Spectrum software (Scanalytix, USA). Individual DC were tracked using the Fiji plug-in MTrackJ (Meijering et al., 2012). Migratory aspects including velocity, distance to starting point and directional persistence were determined using the Chemotaxis and Migration Tool V2.0 (ibidi, Germany).

### Adhesion assay

Adhesion assay was essentially performed as described before (Kurz et al., 2000) with some modifications. 96-well flat bottom plates were coated overnight at 4°C with 100 μg/ml of fibronectin, laminin, and collagen IV (all from Sigma-Aldrich, USA). After washing the coated wells two times with PBS, 1 × 10^5^ cells/well were seeded in duplicates in complete serum-free RPMI 1640 medium. Plates were centrifuged for 1 min at 400 x g and incubated for 45 min at 37°C. Medium supernatant with non-adhered cells was discarded and plates were washed two times with serum-free medium. Adherent cells were detached from wells for analysis by flow cytometry by adding prewarmed 10 mM EDTA/PBS. Before removing the cells from the plates 1 × 10^4^ Dynabeads were added per well to normalize for numbers of detached cells. Retrieved cells with beads were stained for CD11c, MHC class II, langerin/CD207, and EpCAM and analyzed by flow cytometry. Absolute numbers of detached cells were determined and normalized to number of beads.

### Statistics

Statistical analyses were performed using GraphPad Prism 7 software (GraphPad Software Inc., USA). One-tailed unpaired Student’s *t* test was used for pairwise comparisons. For multiple groups comparison one-way ANOVA was performed followed by a post-hoc Tukey’s multiple comparisons test. Differences with a p value < 0.05 were considered statistically significant.

## Results

### Analyses of Met-deficient skin DC under steady state and inflammatory conditions

We have previously demonstrated that Met-signaling impacts on the migration of LC and dDC (dDC) from skin towards draining lymphnodes (dLN) under inflammatory conditions *in vivo* and *in vitro* (Baek et al., 2012). In order to study the mechanism of Met-signaling in DC in more detail, we sought to extend our phenotypic analysis in a conditional Met-KO mouse model. We began with analyzing the DC subsets in Met-KO vs. Met-WT mouse epidermis and dermis under steady-state conditions.

To this end, ear skin of Met-KO and Met-WT mice were split into dorsal and ventral halves. Single cell suspensions from further separated epidermal and dermal sheets were then analyzed with flow cytometry (Fig. 1). The hematopoietic population of the epidermis comprises only γδ T cells commonly known as dendritic epidermal T cells (DETC) and Langerhans cells (LC). LC were identified as CD45+ MHC-II+ CD3− while DETC as CD45+ MHC-II-CD3+ (Fig. 1A). At steady state, no differences were observed in Met-KO vs. Met-WT animals for both LC (Fig. 1B) and DETC (Fig.1C), which indicates that after induction of the Met-KO phenotype no accumulation of Met-deficient LC or DETC occurs. Thus Met-signaling may not play a role in LC and DETC migration in the epidermis at steady state.

**Fig. 1.**
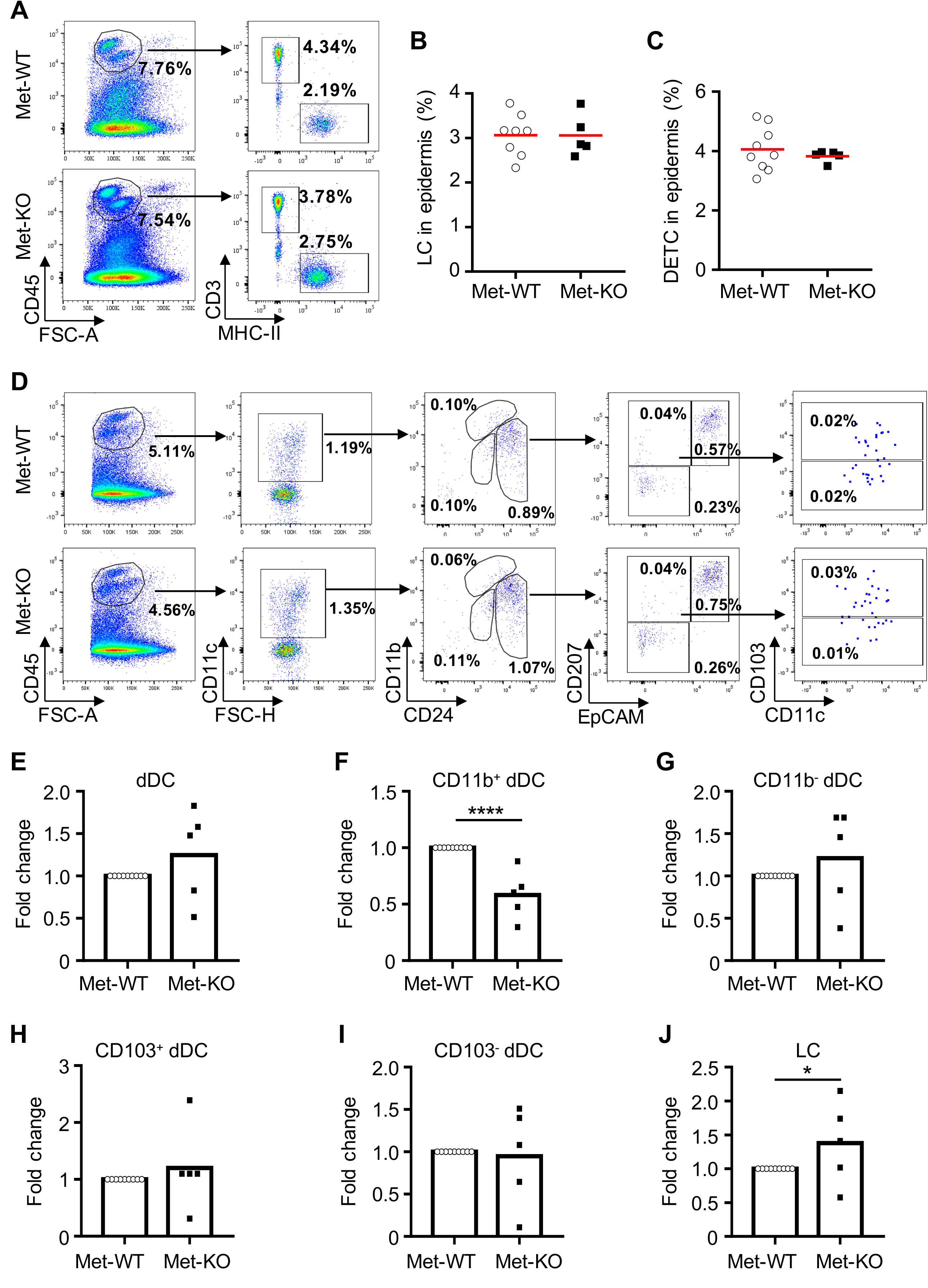
Analysis of LC and dDC subsets in Met-KO and Met-WT skin at steady state. Single cell suspensions from ear epidermal and dermal sheets of Met-KO and Met-WT mice were analyzed by flow cytometry. Events shown in dot plots were pre-gated on live singlet cells. (A) Gating strategy for LC (CD45+ MHC-II+ CD3−) and DETC (CD45+ MHC-II-CD3+) from epidermal single cell suspension of Met-KO and Met-WT skin. (B) LC and (C) DETC percentage in Met-KO and Met-WT epidermis. (D) Gating strategy for dDC (CD11c+), CD11b+dDC (CD11c+ CD11b+ CD24−), CD11b-dDC (CD11c+ CD11b− CD24−), CD103+dDC (CD11c+ CD11b− CD24+ CD207+ CD103+), CD103-dDC (CD11c+ CD11b− CD24+ CD207+ CD103−) and migratory LC (CD11c+ CD11bint CD24+ CD207+ EpCAM+) from dermal single cell suspension of Met-KO and Met-WT skin. (E-J) Bar graphs show (E) dDC, (F) CD11b+dDC, (G) CD11b-dDC, (H) CD103+dDC, (I) CD103-dDC and (J) migratory LC fold change in percentage out of total dermal cells in Met-KO and Met-WT dermis. Percentages of cells from Met-WT were set as 1. Data shown are from three independent experiments (Met-WT, n=8-9; Met-KO, n=5). \**p*<0.05, \*\*\*\**p*<0.0001.

In dermis, five different CD11c+ DC subsets were identified in the hematopoietic compartment (CD45+): (i) CD11b+ dDC, (ii) CD11b− dDC, (iii) CD207+ CD103+ dDC, (IV) CD207+ CD103− dDC and (v) CD207+ EpCAM+ migratory LC (Fig. 1D; Henri et al., 2010). A slight accumulation of CD11c+ DC was observed in the dermis of Met-KO mice compared to the dermis of control mice (Fig. 1E), which can be ascribed to increased numbers of cells of the CD11b− DC compartment. However, only the increase in CD207+ EpCAM+ migratory LC (Fig. 1F-J) was found statistically significant. In contrast, we found a significant decrease of CD11b+ dDC in Met-KO dermis.

Next, the effect of Met-signaling on LC/dDC migration under inflammatory conditions was investigated. Dorsal ear skin halves were isolated and painted with a mixture of acetone and dibutyl phthalate (DBP) to irritate the skin, inducing LC/dDC emigration from *ex vivo* cultured ear tissue into the medium. The cultured dorsal halves were trypsinized after 24 h and 48 h to generate epidermal and dermal single cell suspensions that were analyzed by flow cytometry (Fig. 2).

**Fig. 2.**
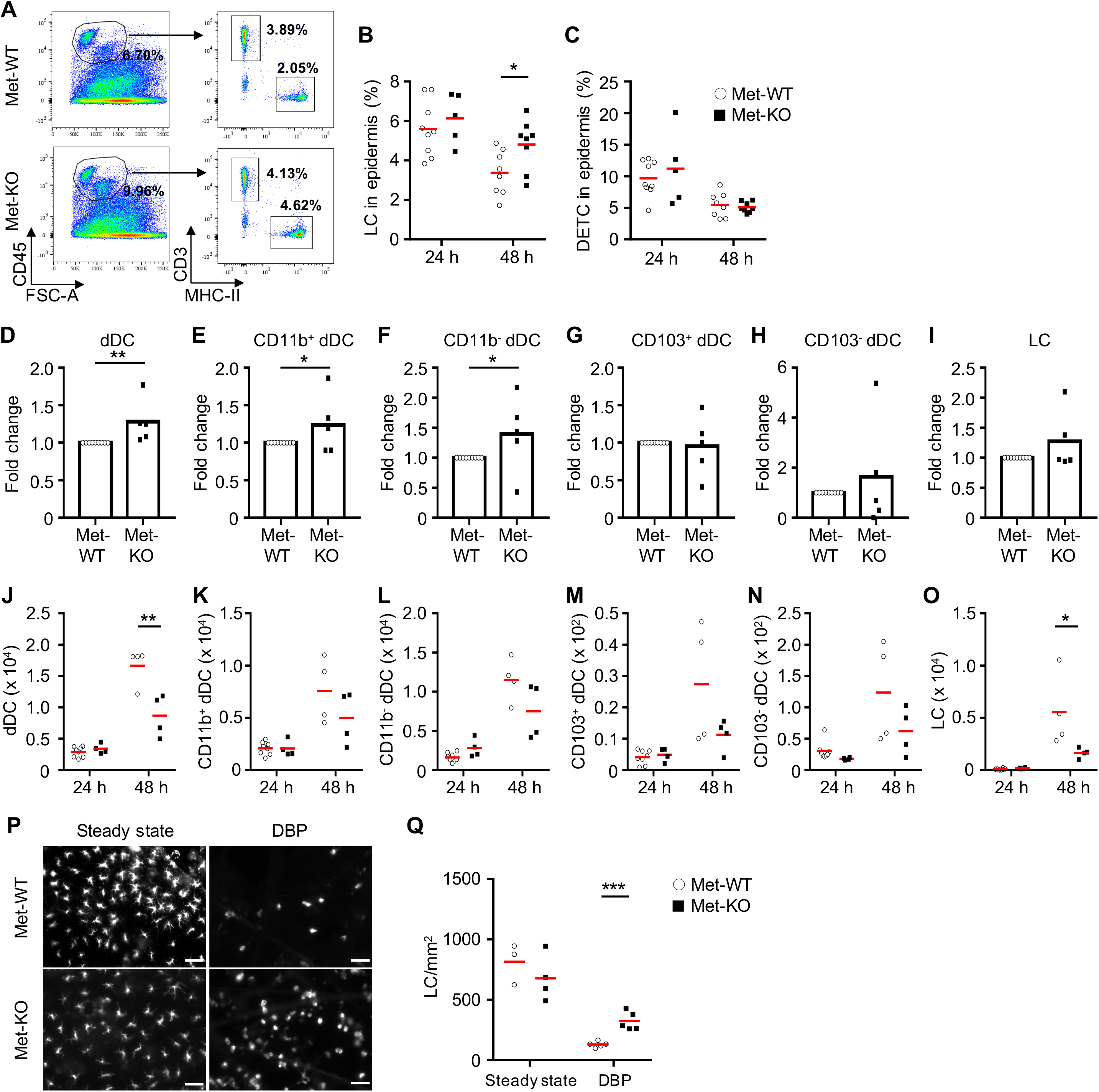
Emigration of LC and dDC subsets from Met-KO and Met-WT skin under inflammatory condition. Ear dorsal halves from Met-KO and Met-WT mice were painted with acetone/DBP and cultured for up to 48 h. (A) Gating strategy for LC (CD45^+^ MHC-II^+^ CD3^−^) and DETC (CD45^+^ MHC-II^−^ CD3^+^) from epidermal single cell suspension of Met-KO and Met-WT skin. Scatter plots show (B) LC and (C) DETC percentage in Met-KO and Met-WT epidermis. Data shown are from three independent experiments (Met-WT, n=8-9; Met-KO, n=5-8). (D-I) Dermal sheets were trypsinized and further treated with collagenase to generate single cell suspension. Gating for (D) dDC, (E) CD11b^+^dDC, (F) CD11b^−^dDC, (G) CD103^+^dDC, (H) CD103^−^dDC and (I) migratory LC was done as shown in Fig. 1. Bar graphs show fold change in percentage out of total dermal cells in Met-KO and Met-WT dermis. Percentages of cells from Met-WT were set as 1. Data shown are from three independent experiments (Met-WT, n=8-9; Met-KO, n=5). (J-O) LC and dDC emigrated into culture medium were analyzed by flow cytometry. Gating for emigrated (J) dDC, (K) CD11b^+^dDC, (L) CD11b^−^dDC, (M) CD103^+^dDC, (N) CD103^−^dDC and (O) LC was done as shown in Suppl. Fig. 1. Scatter plots show the absolute cell count from Met-KO and Met-WT skin explants. Data shown are from three independent experiments (Met-WT, n=4-9; Met-KO, n=4). (P, Q) Isolated epidermal sheets from Met-KO and Met-WT ear skin were fixed in cold acetone at steady state or after painting with acetone/DBP and culturing for 48 h, and stained with anti-MHC-II to identify LC. (P) Representative microscopic images of LC in epidermal sheet. Scale bar, 50 μm. (Q) Scatter plot shows LC count/mm^2^. Data shown are from three independent experiments (Met-WT, n=3-5; Met-KO, n=4-5). \**p*<0.05, \*\**p*<0.01, \*\*\**p*<0.001.

In Met-KO epidermis accumulation of LC was observed. After 48 h, the percentage of LC remaining in Met-KO epidermis was significantly higher than that of the Met-WT control, indicating that Met-KO LC cannot efficiently migrate out of the epidermis (Fig. 2A, B). This is in line with our previous findings (Baek et al., 2012). In contrast, the percentage of DETC significantly decreased over time in both Met-KO and Met-WT epidermis, thereby showing that Met-signaling does not play a role in DETC emigration from skin (Fig. 2C).

In the dermis, Met-KO dDC migration was remarkably impaired compared to that of Met-WT dDC, leading to the accumulation Met-KO dDC in the dermis after 24 h (Fig. 2D-I). However, only the CD11b+ and CD11b− dDC subsets showed significantly impaired migration.

Furthermore, we analyzed the medium for emigrated LC/dDC after 48 h by flow cytometry (Fig. 2J-O and Suppl. Fig. 1A). Since retention of Met-KO LC and certain dDC subsets was observed in the epidermis and dermis respectively, less emigrated Met-KO LC/dDC were expected in the medium compared to the Met-WT counterpart. Indeed, higher numbers of dDC emigrated from Met-WT skin compared to the Met-KO skin (Fig. 2J). All Met-KO dDC subsets showed decreased emigration, however only Met-KO LC emigration was found to be statistically significant (Fig. 2K-O).

This observation was further corroborated by immunofluorescence analysis of epidermal sheets from Met-KO and Met-WT skin 48 h after DBP treatment. In line with the flow cytometry analyses shown before (Fig. 2A, B) a significantly higher LC density was observed in Met-KO epidermis than in the Met-WT counterpart (Fig. 2P, Q).

Taken together, these data demonstrate that Met-signaling impacts on the migratory properties of both epidermal LC and dDC in skin under inflammatory conditions but leaves migration of DETC unaffected. Our data further suggest that Met-signaling could also play a role in dDC migration in steady state.

### The role of Met-signaling in LC detachment and mobilization

LC are integrated within the layer of keratinocytes in the epidermis through various cell-cell adhesion molecules, including E-cadherin and EpCAM (Tang et al., 1993; Borkowski et al., 1996; Alvarez et al., 2008). LC maturation is associated with the downregulation of E-cadherin and EpCAM, thereby detaching LC and facilitating their migration (Schwarzenberger and Udey, 1996; Bobr et al., 2012; Kel et al., 2012; Brand et al., 2020).

Since Met-deficient LC fail to emigrate efficiently out of the epidermis, we hypothesized that Met-signaling may be necessary for the loss of cell-to-cell contacts by adhesion molecules in LC upon their maturation. Therefore, the expression of E-cadherin and EpCAM was analyzed in Met-KO and Met-WT LC by flow cytometry. Epidermis from Met-KO and Met-WT skin explants were either freshly isolated and trypsinized immediately or painted with acetone/DBP and cultured for 24 h and 48 h before isolation and trypsinization (Fig. 3). Both Met-KO and Met-WT LC were found to downregulate the surface expression of E-cadherin and EpCAM after painting with acetone/DBP (Fig. 3A, B), which was further supported by expression data on the transcriptional level of sorted Met-KO and Met-WT LC using qPCR (Suppl. Fig. 1B). This indicates that Met-signaling does not play a role in LC detachment.

**Fig. 3.**
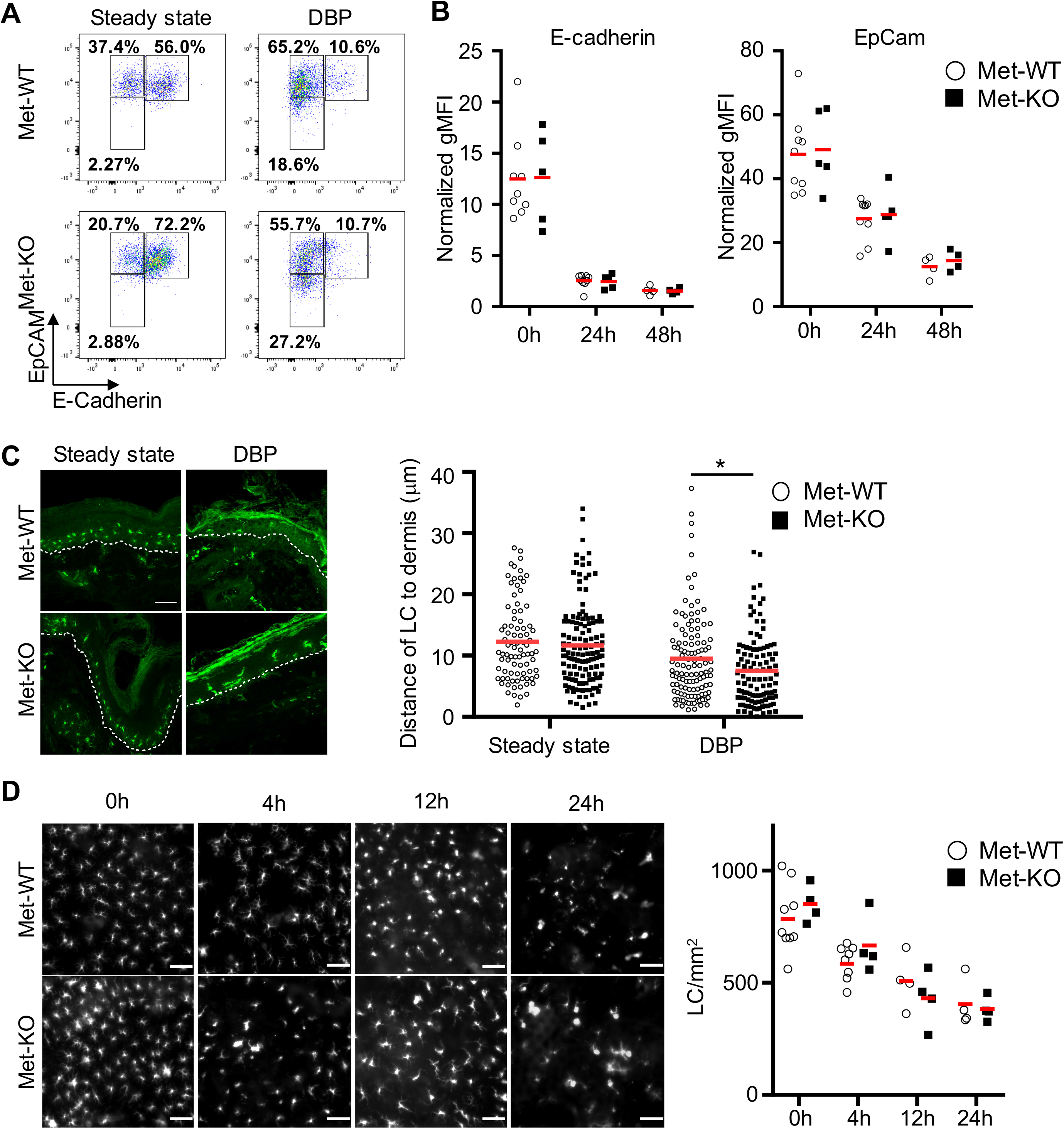
Migratory capacity of LC within Met-KO and Met-WT epidermis. (A, B) Expression of E-cadherin and EpCAM on LC at steady state and after painting with acetone/DBP and culturing for 24 h and 48 h was analyzed by flow cytometry. Events shown in dot plots were pre-gated on live singlet cells, CD45+ and MHC-II+. Scatter plots show normalized gMFI of E-cadherin (left panel) and EpCAM (right panel) at steady state and after culture. Data shown are from three independent experiments (Met-WT, n=4-9; Met-KO, n=4-5). (C) Footpad skin was isolated from Met-KO and Met-WT mice and cryopreserved in OCT either before or after painting with acetone/DBP and culturing for 24 h. Then, the skin was sectioned and stained with anti-MHC-II and anti-CD49f Abs and DAPI. Representative microscopic images of LC and dDC (green) in skin are shown (left panel). Dotted line represents the epidermal dermal junction (EDJ). Scale bar, 50 μm. Scatter plot shows the distance of LC to EDJ at steady state and after acetone/DBP painting (right panel). At least 87 cells were analyzed per condition. Data shown are from two independent experiments with n=2-4 mice per group. (D) Basement membrane depleted epidermal sheets from Met-KO and Met-WT ear skin were isolated at steady state. Isolated epidermal sheets were either fixed in cold acetone immediately or after culture for 4 h, 12 h or 24 h and stained with anti-MHC-II Ab. The left panel shows representative microscopic images of LC in epidermal sheet. Scale bar, 50 μm. Scatter plot (right panel) shows LC count/mm^2^. Data shown are from three independent experiments (Met-WT, n=4-9; Met-KO, n=4-5). \**p*<0.05.

We thus addressed next, whether Met-deficient LC have a mobilization defect that retains them within the epidermis or fail to degrade and cross the basement membrane. To this end, skin explants from Met-KO and Met-WT mice were isolated in steady state or 24 h after painting with aceton/DBP. Skin sections were stained with anti-MHC-II and anti-CD49f antibodies and analyzed by confocal fluorescence microscopy to visualize the longitudinal distribution of LC in the epidermis and the epidermal dermal junction (EDJ), respectively.

In steady state, Met-KO and Met-WT LC are similarly distributed in the epidermis (Fig. 3C and Suppl. Fig. 2). Upon activation, emigration of LC is induced, however more Met-KO LC are retained in the epidermis compared to Met-WT LC (Fig. 3C). Notably, the retained Met-KO LC were found lined across the EDJ, while Met-WT LC were more sparsely distributed (Fig. 3C). Next, the distance of individual Met-KO and Met-WT LC to the EDJ was determined at steady state and under inflammatory conditions. LC from both Met-KO and Met-WT epidermis were found closer to the EDJ after painting with acetone/DBP, as the average distance of individual Met-KO and Met-WT LC to the EDJ decreased from ~12 μm at steady state conditions to ~8 μm 24 h after the treatment (Fig. 3C). This shows that, similar to Met-WT LC, Met-KO LC are mobile and capable of moving towards the basement membrane. However, the basement membrane appears to constitute a physical barrier that impedes the transmigration of Met-KO LC to the dermis, leading to the retention of Met-KO LC. This could also explain the fact that the average distance of Met-KO LC to the EDJ was significantly less than that of Met-WT LC, which remained in the epidermis (Fig. 3C).

For this reason, we next investigated LC emigration from the epidermis in the absence of the basement membrane. To this end, we separated the epidermis from the dermis at exactly the upper lamina lucid of the basement membrane using EDTA, leaving the basement membrane attached to the dermis (Zou and Maibach, 2018). Epidermal sheets from Met-KO and Met-WT ear skin were stained for MHC-II at steady state or after 4 h, 12 h, and 24 h of culture (Fig. 3D). As LC emigrate from the epidermis, LC density was dramatically reduced in both Met-KO and Met-WT epidermal sheets after culture. Moreover, there was no significant retention of Met-KO LC in the epidermis compared to Met-WT LC (Fig. 3D). This shows that the Met-KO LC can efficiently migrate out of the epidermis in the absence of the basement membrane.

Taken together, our data revealed that Met-signaling is neither essentially involved in LC detachment from surrounding tissue nor in the initiation of LC motility and emigration. Importantly, our results also shows that Met-signaling is critically involved in the ability of LC to cross the basement membrane.

### Met-signaling impacts on podosome formation in DC that regulates ECM degradation

The basement membrane between epidermis and dermis forms a continuous layer suggesting that LC need to degrade it to pass through and reach the dermis. A study by Stoitzner et al. has shown that LC extend protrusions to penetrate the basement membrane demonstrating the absence of intact basement membrane surrounding these pseudopods (Stoitzner et al., 2002). These protrusions resemble podosomes, which are highly dynamic actin-based structures that are known to play a role in the invasiveness of macrophages and DC by coordinating cytoskeleton rearrangement and MMP activity leading to ECM degradation (Gawden-Bone et al., 2010; Murphy and Courtneidge, 2011; van den Dries et al., 2014; Wiesner et al., 2014). Moreover, it was found that in some invasive tumors podosome formation was dependent on Met-signaling (Rajadurai et al., 2012). This raised the question whether the formation of podosomes in DC is dependent on Met.

Therefore, we next examined the capacity of Met-KO DC to form podosomes and degrade the ECM. For this, BMDC were incubated overnight on coverslips coated with a thin layer of Oregon green-labelled gelatin. During the incubation period, BMDC form podosomes and degrade the fluorescent gelatin, creating non-fluorescent patches. BMDC were then fixed and the actin cytoskeleton was stained with phalloidin to visualize podosome forming cells (Fig. 4A). The immunofluorescence imaging revealed that podosome clusters of DC colocalize with degraded regions. The total area of the degraded gelatin patches was quantified. Interestingly, Met-KO BMDC showed significantly lower ability to degrade the gelatin compared to the Met-WT counterpart (Fig. 4B). Consistently, the percentage of Met-KO BMDC forming podosomes was 2-fold lower than that of the Met-WT BMDC (Fig. 4C). To ensure that Met-KO BMDC do not suffer from an adhesion defect that prevents them from efficiently forming podosomes and degrading the gelatin, the ventral cell area of Met-KO and Met-WT BMDC was determined, however, no significant differences were observed (Fig. 4D). No difference was also observed for the distribution of Met surface expression, which did not exhibit a specifically enhanced colocalization with podosome clusters (Suppl. Fig. 3A). We next quantified the number of actin cores within podosome clusters in Met-KO and Met-WT BMDC. Although the number of actin cores varied greatly ranging from 20 to 260 cores per cell, Met-deficient BMDC showed remarkably less actin cores per cell compared to the WT control BMDC (Fig. 4E, F). However, co-staining of BMDC for vinculin, a component of the ring structure surrounding the actin cores, revealed that the integrity of podosome clusters is not affected by Met deficiency (Suppl. Fig. 3B). We also found no evidence for a different transcriptional regulation of expressed MMPs (MMP2, MMP9 and MT1MMP/MMP14; Suppl. Fig. 1C) that are well recognized as podosome associated proteases involved in ECM degradation (Murphy and Courtneidge, 2011; Wiesner et al., 2014).

**Fig. 4.**
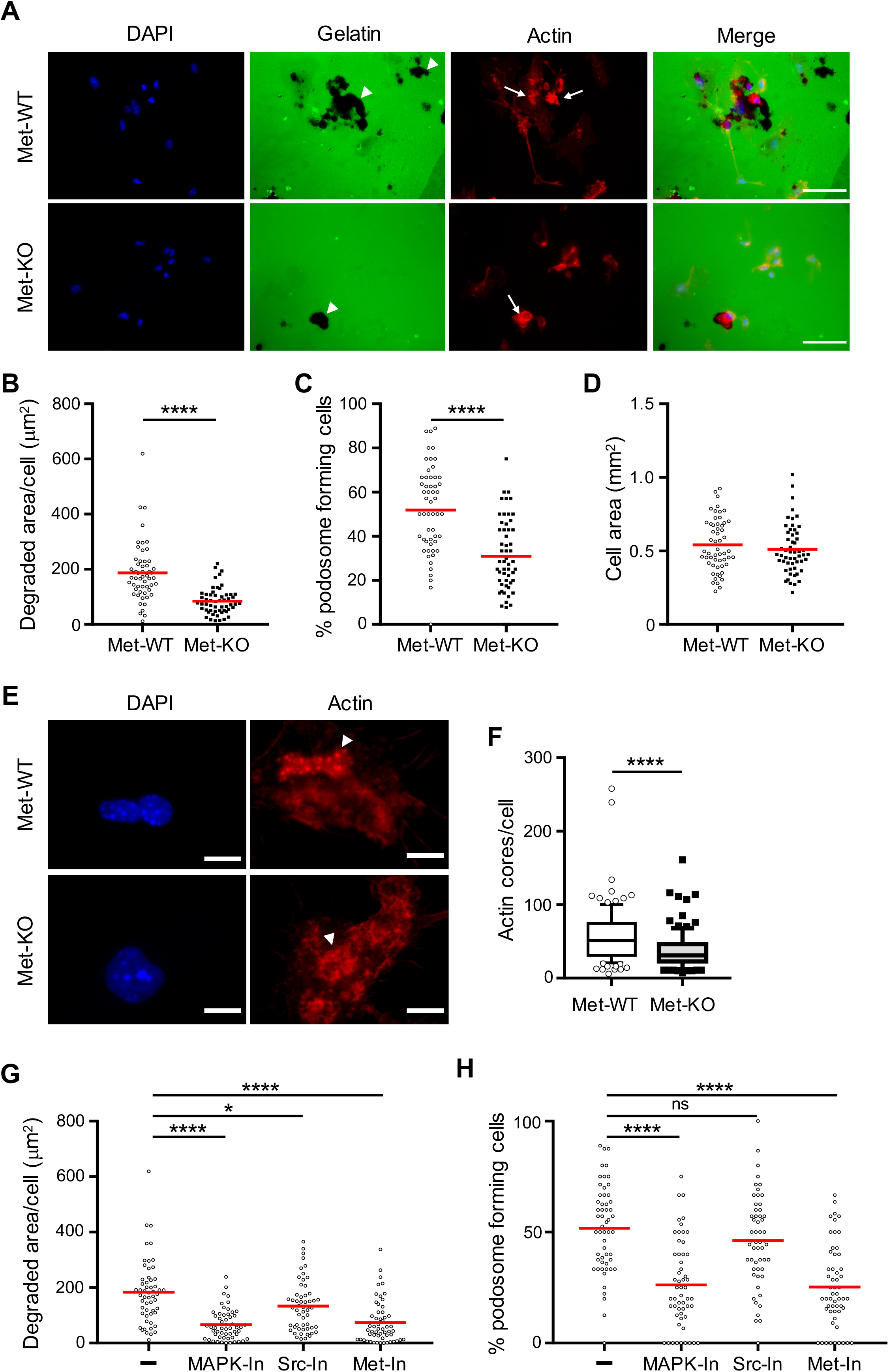
Met-signaling deficiency impairs podosome formation in DCs and ECM degradation. (A) Immunofluorescence microscopy of BMDC from Met-WT and Met-KO mice cultured on fluorescently labeled gelatin (green) for 24 h. Arrowheads indicate degraded areas, arrows point at podosomes. Scale bar, 50 μm. (B-D) Diagrams show (B) size of degraded area per cell, (C) fraction of DCs that were found podosome positive, and (D) size of the attached cell bodies. Data are from three independent experiments with a total of 55 sections analyzed for each condition. (E) Immunofluorescence microscopy of BMDCs from Met-WT and Met-KO at higher magnification for visualization of actin cores within podosomes (arrowheads). Scale bar, 10 μm. (F) Number of actin cores per cell. Box-whisker plot shows mean values ± 25th and 75th percentile (boxes) and 10th and 90th percentile (whiskers), respectively (n=3 with ≥ 30 cells per experiment and condition). (G, H) Gelatin degradation and podosome formation assays were performed with Met-WT BMDC treated with and without 5 μM PD98059 (MAPK-In), 2 μM SU6656 (Src-In) or 1 μM SU11274 (Met-In). Scatter plots show (G) degraded area and (H) percentage of podosome forming cells. Data are from three independent experiments with at least 15 fields analyzed per experiment; ns, not significant; \**p*<0.05, \*\*\*\**p*<0.0001.

The activation of the Met-signaling pathway leads to the recruitment of various downstream signaling adaptor molecules, such as Grb2 and Gab1, which serves as docking platforms to activate various downstream signaling axes, including the Ras-MEK-MAPK/ERK and the Src-STAT3 signaling pathways (Trusolino et al., 2010; Gherardi et al., 2012; Ilangumaran et al., 2016;). Similarly, we described before that upon HGF treatment mature BMDC exhibited fast and sustained activation of ERK1/2 (Baek et al., 2012). Therefore, we employed the MEK inhibitor PD98059 (MAPK-In), the Src kinase inhibitor SU6656 (Src-In) and the Met kinase inhibitor SU11274 (Met-In) to dissect the Met-signaling pathway and identify which signaling axis is crucial for the Met-dependent podosome formation and the consequent gelatin degradation in BMDC. Met-WT BMDC were applied on fluorescent gelatin-coated coverslips in the presence of TNFα and with or without inhibitors. Following overnight incubation, the gelatin degradation and podosome formation were assessed (Fig. 4G, H and Suppl. Fig. 4).

BMDC treated with the Met-In and the MAPK-In showed a 2-fold decrease in the degraded area per cell when compared to the untreated BMDC, while Src-In treated BMDC showed only a slight decrease (Fig. 4G). In line with this, only Met-In and MAPK-In treated BMDC, but not Src-In treated BMDC, showed a significant reduction in the percentage of podosome-forming cells compared with untreated BMDC. (Fig. 4H). In contrast, treatment with the aforementioned inhibitors had no adverse effects on the ventral cell area of BMDC (Suppl. Fig. 4B). Notably, the observed reduction in degradation activity and podosome formation was similar in both the Met-In and the MAPK-In treated BMDC to that observed in the Met-KO BMDC (Fig. 4B).

Taken together, this results demonstrate that Met-signaling is critically involved in podosome formation and function, that regulate MMP activities leading to ECM degradation. This is in line with our previous report on the regulation of MMP activities by Met-signaling in DC (Baek et al., 2012). Moreover, the data provide strong evidence that the MAPK/ERK signaling axis is indispensable for Met-dependent podosome formation and ECM degradation.

### Met-signaling impacts on motility but not chemotaxis of DC in 3D collagen gel migration

Met deficiency impairs the migration of not only LC, but also dDC *ex vivo* and *in vivo* (Fig. 2) (Baek et al, 2012). However, dDC, unlike LC, do not need to cross a continuous dense basement membrane separating the epidermis and dermis. In the dermis, LC and dDC migration to the sdLN is dependent on CCR7 and its respective ligands: CCL19 and CCL21. It is worth mentioning that LC migration from the epidermis to the dermis is CXCR4-dependent but CCR7-independent as observed in CCR7-deficient mice (Ohl et al., 2004). Additionally, studies from the Sixt lab have indicated that DC migration in the dermal compartment is independent of ECM remodeling and pericellular proteolysis. Instead, DC migrate by squeezing and flowing through the path of the least resistance guided by the CCL19/CCL21 gradient until they reach the initial lymph vessel. There they again squeeze themselves through the discontinuous basement membrane surrounding the initial lymph vessel into the lumen, thus, questioning the role of podosomes for interstitial migration of DC (Lämmermann et al., 2008; Renkawitz et al., 2019).

To mimic DC migration through the extracellular matrix-rich dermal interstitial space, we employed a protocol that was established by Sixt et al. in which chemotaxis of DC is visualized *in vitro* in 3D collagen gels by time-lapse bright-field microscopy (Sixt and Lämmermann, 2011). BMDC were generated from Met-KO and Met-WT BM and treated with TNFα overnight to induce their maturation. Then, BMDC were embedded in collagen and their migration through the collagen gel was visualized for 4 hours in the presence and absence of a CCL19 chemokine gradient. Trajectory plots showed that Met-deficient BMDC, similar to WT control BMDC, respond to the CCL19 gradient and migrate towards it (Fig. 5A-D). Analysis of migratory properties including velocity, eucledian distance and directional persistence clearly showed a significant increase in all these parameters in the presence of the CCL19 gradient, however no differences were observed between Met-KO and Met-WT BMDC (Fig. 5B-D). In line with this, we found no evidence for a different transcriptional regulation of CCR7 and CXCR4 expression in Met-KO and Met-WT LC (Suppl. Fig. 1D). These data confirm and extend our previous observation of DC migration using Boyden chamber assay (Baek et al., 2012) that chemokine receptor-mediated chemotactic response is not affected by Met-deficiency.

**Fig. 5.**
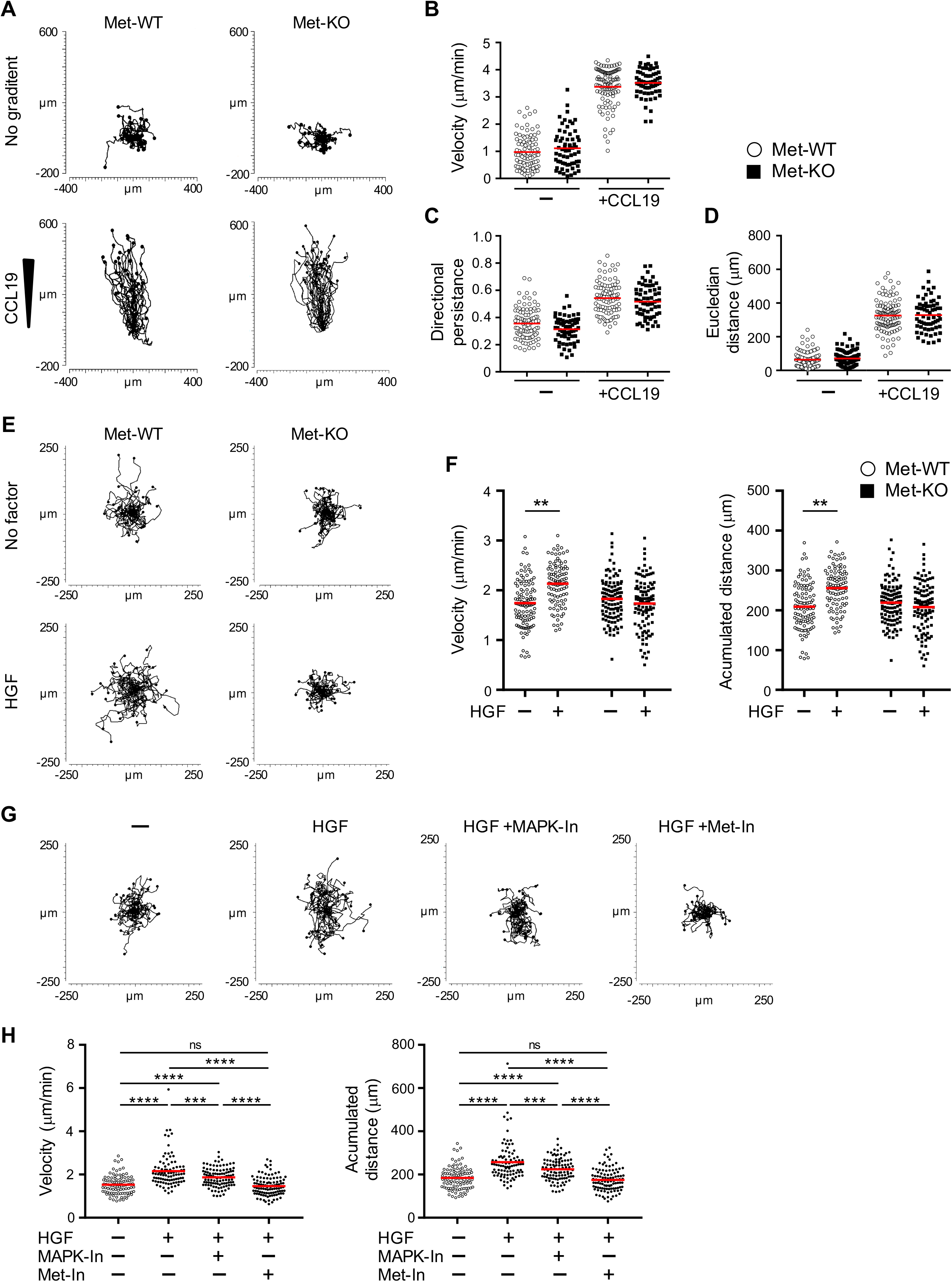
Impact of Met-signaling on motility and chemotaxis of DC in 3D collagen gel migration. TNFα treated BMDC were embedded in collagen gels and their migration was visualized using time-lapse bright field microscopy. Data were pooled from three independent experiments with ≥ 30 cells for each condition per experiment. (A-D) Chemotactic response of mature BMDC in the absence and presence of CCL19 gradient was tracked for 4 h. (A) Representative trajectory blots of Met-KO and Met-WT BMDC. Each dot represents one cell. Scatter plots show (B) velocity, (C) directional persistence and (D) eucledian distance of Met-KO and Met-WT BMDC in the absence and presence of CCL19 gradient. (E, F) Motility of mature BMDC in response to HGF treatment. Met-KO and Met-WT BMDC were treated with TNFα overnight in the presence or absence of HGF. Migration of embedded cells was tracked for 2 h. (E) Representative trajectory blots of Met-KO and Met-WT BMDC with or without HGF treatment. Each dot represents one cell. (F) Scatter plots show velocity (left panel) and accumulated distance from starting point (right panel) of Met-KO and Met-WT BMDC with or without HGF treatment. (G, H) Impact of Met-signaling on motility of mature BMDC. Mature Met-WT BMDC were treated with HGF in the presence or absence of 5 μm PD98059 (MAPK-In) or 1 μm SU11274 (Met-In). Migration of embedded cells was tracked for 2 h. (G) Representative trajectory blots of BMDC in absence and presence of HGF with or without MAPK-In and Met-In. Each dot represents one cell. (H) Scatter plots show velocity (left panel) and accumulated distance from starting point (right panel); ns, not significant; \*\**p*<0.01, \*\*\**p*<0.001, \*\*\*\**p*<0.0001.

However, Ólafsson et al. demonstrated recently that HGF indeed induces motility of BMDC within collagen gels in a MAPK/ERK-dependent manner (Ólafsson et al., 2020). Therefore, we examined HGF-induced motility of Met-KO and Met-WT BMDC in 3D collagen gels. BMDC were stimulated with TNFα or left untreated in the presence or absence of HGF. BMDC were then embedded in collagen and their motility within the collagen gel was recorded for 2 hours. Trajectory plots of immature Met-KO and Met-WT BMDC were similar in the presence and absence of HGF and no differences were observed for their velocity and accumulated distance traveled (Suppl. Fig. 5). In contrast, Met-WT BMDC upon maturation with TNFα exhibited increased motility in response to HGF, resulting in increased velocity and accumulated distance traveled, which was not observed in TNFα treated Met-KO BMDC (Fig. 5E, F).

Furthermore, we assessed the importance of the MAPK/ERK signaling axis in HGF-induced motility in BMDC by employing specific inhibitors (Fig. 5G, H). TNFα matured Met-WT BMDC were treated with HGF in the presence or absence of MAPK-In and Met-In. Trajectory plots demonstrated that HGF-induced motility is abolished in the presence of either the MAPK-In or Met-In (Fig. 5G). Both inhibitors significantly diminished the HGF-induced increase in BMDC velocity and accumulated distance traveled (Fig. 5H). These results further corroborates the important role of the MAPK/ERK signaling pathway in Met-signaling mediated DC motility and migration regulation.

### Met-signaling regulates adhesion of LC to ECM factors

After crossing the epidermal basement membrane, LC have to migrate through the ECM-rich dermis. Depending on the architecture and topology of ECM-rich structures, LC alternate between adhesion-dependent and adhesion-independent migration (Loh and Su, 2016). Further studies have demonstrated that HGF modulates the adhesive properties of DC and other immune cells (Adams et al., 1994; Mine et al., 1998; Skibinski et al., 2001; Kurz et al., 2002). Therefore, we sought to determine the effect of HGF on the adhesion of BM-derived LC from Met-KO and Met-WT mice to different components of the extracellular matrix, namely fibronectin, laminin and collagen IV. BMLC were generated from Met-KO and Met-WT BM as described previously and treated with HGF overnight or left untreated (Capucha et al., 2018). No impact was observed on the development of BMLC in culture due to the Met-deficiency (Suppl. Fig. 6) and in line with previous findings for LC/DC development *in vivo* and *in vitro* (Fig. 1; Baek et al. 2012).

**Fig. 6.**
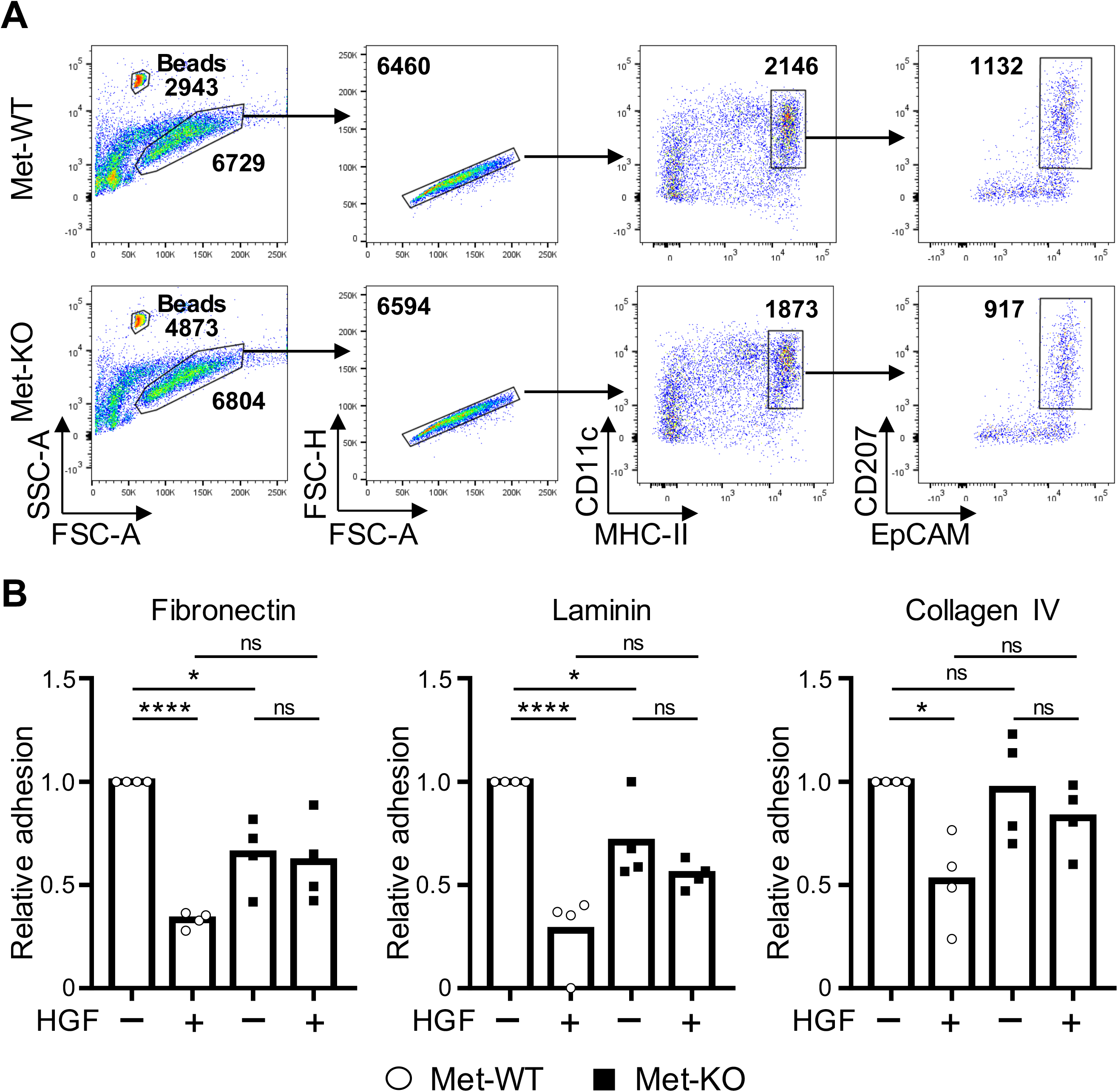
Met-signaling impacts on BMLC adhesion to ECM components. TGF-β-derived BMLC were generated from Met-KO and Met-WT BM and treated with or without HGF overnight. Then, BMLC adhesion to ECM factors fibronectin, laminin and collagen IV was assessed. (A) Representative flow cytometry plots of the analysis of adhered BMLC (CD11c^+^ MHC-II^+^ CD207^+^ EpCAM^+^) to wells coated with fibronectin showing the gating strategy after detachment from ECM factors. Absolute numbers of beads and cells are shown. (B) Bar graphs show the relative adhesion of Met-KO and Met-WT BMLC to fibronectin (left panel), laminin (middle panel), and collagen IV (right panel). Absolute count of Met-WT BMLC was set as 1. Data shown are from two independent experiments, each with two technical replicates; ns, not significant; \**p*<0.05, \*\**p*<0.01, \*\*\**p*<0.001, \*\*\*\**p*<0.0001.

BMLC were allowed to adhere to the various ECM components and non-adherent cells were discarded. After adhered cells were detached and recovered, BMLC were analyzed and quantified using flow cytometry (Fig. 6 and Suppl. Fig. 6A). Met-WT BMLC treated with HGF exhibited a reduced adhesion to fibronectin, laminin and collagen IV in comparison to untreated BMLC, suggesting that Met-signaling by HGF reduces BMLC adhesion to the ECM. In contrast, HGF treatment had no impact on Met-deficient BMLC, as both untreated and HGF treated Met-KO BMLC showed a similar adhesion to fibronectin, laminin and collagen IV. In general, Met-KO BMLCs showed lower adhesion to fibronectin and laminin than untreated Met-WT BMLCs, while adhesion to collagen did not differ significantly. This is in accordance to the results obtained in the motility assay in 3D collagen gels (Fig. 5E,F). Accordingly, it can be assumed that the enhanced motility of Met-WT DC in 3D collagen gels upon HGF treatment, which is not found with Met-deficient DC, is due to attenuated adhesion following HGF-mediated Met-signaling. Taken together, our data clearly indicate that Met-signaling impact on both adhesion-dependent and adhesion-independent migration of LC/DC.

## Discussion

It became increasingly evident that the HGF/Met-signaling pathway is involved in the regulation of immune function at least in part by regulating motile and migratory properties of immune cells, including DC (Molnarfi et al., 2015; Ilangumaran et al., 2016; Sagi and Hieronymus 2018). By employing a conditional Met knockout mouse model, we demonstrate in this study that Met-signaling is critically involved in podosome formation in DC and thus in their capacity to efficiently degrade ECM factors. This appears as an important mechanism for LC to penetrate the continuous epidermal basement membrane for their migration through the dermis towards dLN, while their detachment from the surrounding tissue and chemotactic response were not evidently affected by Met deficiency.

LC development and migration at steady state are not dependent on Met-signaling, however, Met-deficient LC accumulate in epidermis during inflammatory conditions (Baek et al., 2012). At steady state, LC express various epithelial markers, including E-cadherin and EpCAM, which are necessary for the interaction of LC with keratinocytes, thereby integrating LC functionally within the epidermis, rendering LC sessile (Tang et al., 1993; Borkowski et al., 1996; Sagi and Hieronymus, 2018). Here, we describe that LC downregulate the expression of these epithelial markers upon maturation in agreement with other studies (Schwarzenberger et al., 1996; Jiang et al., 2007, Bobr et al., 2012; Kel et al., 2012). However, we found that the expression regulation of E-cadherin and EpCAM is not dependent on Met-signaling. Surprisingly, a recent study by Brand et al. showed that specific deletion of E-cadherin in CD11c+ cells in mice had no effect on LC anchorage in the epidermis (Brand et al., 2019). In another study, deletion of EpCAM in langerin+ cells in mice had also no effect on LC anchorage, on the contrary, LC could not migrate out of the epidermis in these mice (Gaiser et al., 2012). This shows that the LC anchorage process is complex and not yet fully understood. Moreover, we found here that Met-deficient LC move towards the dermis but line up over the basement membrane, failing to penetrate it. However, in the absence of the basement membrane Met-deficient LC can migrate efficiently out of the epidermis, which supports the notion that the Met-deficient LC cannot degrade the basement membrane.

LC penetrate the basement membrane through the formation of protrusive cytoplasmic structures that resemble podosomes (Stoitzner et al., 2002), which are known to play a role in ECM remodeling by coordinating the cytoskeleton rearrangement and MMP activity, thus promoting invasiveness of DC (Gawden-Bone et al., 2010; Murphy and Courtneidge, 2011; Linder and Wiesner, 2015). Mature DC express various MMP to degrade ECM and basement membrane barriers during their migration. For instance, MMP2 and MMP9 were deemed crucial for the *ex vivo* and *in vivo* migration of LC and dDC (Ratzinger et al., 2002; Yen et al., 2008, Baek et al., 2012). Podosomes recruit three main MMP, namely MMP2, MMP9 and MMP14 (Murphy and Courtneidge, 2011; Wiesner et al, 2014). In DC, podosome-mediated ECM degradation is dependent only on MMP14, as deletion of MMP14, but not of MMP2 or MMP9, abolished podosome-mediated ECM degradation (West et al., 2008; Gawden-Bone et al., 2010). Despite that, MMP2, MMP9 and MMP14 deficiency had no impact on neither the frequency of DC forming podosomes nor the podosome structural integrity (West et al., 2008). In line with this, we found that Met deficiency had no effect on the transcriptional regulation of MMP2, MMP9 or MMP14 in LC and GM-CSF-derived BMDC upon maturation (Baek et al., 2012, and data not shown). This suggests that Met-signaling could be involved in regulating podosome formation kinetic i.e. the assembly and/or disassembly of podosome cores, presumably through the downstream MAPK/ERK signaling axis (Fig. 4).

Results of our previous DC migration studies in Met-deficient mice *in vivo* provided strong evidence that, in addition to LC, dermal DC migration is also regulated by Met-signaling (Baek et al., 2012). Here, we extended our analysis to all DC subsets known to be present in dermis. We observed a remarkably impaired migration of Met-deficient dDC under inflammatory conditions compared to Met-WT dDC, leading to the accumulation of Met-KO dDC in the dermis after 24 hours. Interestingly, not all Met-KO dDC subsets showed significantly impaired migration but only the Langerin-EpCAM-CD11b+ and CD11b− dDC subsets, which account for around 80% of the DC population in dermis (Henri et al., 2010). This suggests that the impact of Met-signaling on different dDC subsets may not be the same but may even be regulated by different migration mechanisms. Indeed, different migration modes have been described for DC migration. Studies by Lämmermann et al., 2008, for example, have shown that DC migration can occur integrin-independent, as DC migrate by flowing and squeezing, which is known as amoeboid migration. The amoeboid migration mode is characterized by weak adhesion to ECM and the absence of pericellular proteolysis. Instead, DC manage to find and migrate along the path with the least resistance, which has been recently shown by Renkawitz et al. for DC migration through 3D fibril collagen (Renkawitz et al., 2019). However, these findings are not consistent with many other studies that have described an indispensable role of MMP, especially MMP9, in the migration of DC in the skin, which suggests that a certain degree of ECM remodeling is necessary for efficient dDC migration (Kobayashi et al., 1999; Ratzinger et al., 2002; Yen et al., 2008; He et al., 2018). Loh et al. has suggested that DC locomotion is highly plastic in nature, switching constantly between adhesion-dependent (mesenchymal) and adhesion-independent (amoeboid) migration modes (Loh and Su, 2016). In line with this, several studies have shown that podosome formation in DC is dictated by environmental cues, including the architecture and topology of the environment, rather than their maturation status (Cougoule et al., 2018; van den Dries et al., 2012). Moreover, it has been clearly demonstrated for macrophages that they can switch between these two modes of migration, which is accompanied by a change in podosome structure (described as 2D and 3D podosomes) (Wiesner et al., 2014). Therefore, we assume that Met-regulated podosome formation may indeed also play a role in dDC migration. Intriguingly, the interplay of components of the podosome architecture on the nanoscale using super-resolution microscopy was just recently described for human DC (van den Dries et al., 2019). It will be interesting to determine how Met-signaling interacts with these factors.

In this study, we show that Met-induced regulation of podosome formation in DC is ligand-independent, as these experiment were done in the absence of HGF. It is consistent with our other results demonstrating that HGF does not only promote BMDC motility in 3D environment but also reduces BMLC adhesion to different ECM components. It is tempting to speculate that HGF-triggered activation of Met promotes adhesion-independent amoeboid migration, while activation of Met in absence of HGF promotes adhesion-dependent, podosome-driven mesenchymal migration in DC.

Various studies have shed the light on Met crosstalk, which revealed that Met-signaling can be regulated in HGF-dependent and HGF-independent manners by a large variety of Met-interacting molecules. These molecules include plexins, semaphorins, other RTK, and integrins (Trusolino et al., 2010; Viticchiè and Muller, 2015). Met-integrins interaction are potentially necessary for podosome formation, as integrins are crucial components of podosomes. Ablation of β1, β2 and β3 integrins eliminated podosomes in osteoclasts (Schmidt et al., 2011). Meanwhile, in vitro-generated BMDC and freshly isolated DC from β2-integrin-null mice failed to form podosomes, indicating that β2 integrins are essential for formation of podosomes in DC (Gawden-Bone et al., 2014). One of the most studied Met-integrin interaction is the association of Met with the β1 subunit of the fibronectin receptor α5β1 integrin. The HGF-independent activation of Met by the β1 integrin has shown to promote the invasion and metastasis in different types of cancer (Jahangiri et al., 2017; Ju and Zhou, 2013; Mitra et al., 2011). Interestingly, Van Helden et al. found human DC podosomes to be enriched in active β1 integrin. They further show that maturation-induced podosome dissolution was associated with a loss in β1 integrin activity, while the use of β1 integrin activating antibody led to impaired maturation-induced podosome dissolution (van Helden et al., 2006). This may suggest that the Met/β1 integrin interaction and the consequent Met activation is essential for MAPK/ERK-dependent podosome formation in DC. Hence, this may represent a possible mechanism for the observed impaired podosome formation in Met-signaling deficient DC in the absence of HGF.

Obviously, the physiological activities commenced by Met-signaling in DC and other immune cells suggest, on the one hand, that the HGF/Met pathway could be a potential target for the treatment of inflammatory diseases, autoimmune disorders, and transplantation (Molnarfi et al., 2015; Ilangumaran et al., 2016). On the other hand, aberrant Met-signaling and overexpression of HGF have been associated with tumor progression and metastasis, thus the Met RTK is emerging as a promising therapeutic target for cancer treatment (Trusolino et al., 2010; Gherardi et al., 2012, Garon and Brodrick, 2021). Remarkably, it was reported that specific Met ablation in neutrophils, impaired their recruitment to the tumor tissue, thus enhancing tumor growth and metastasis (Finisguerra et al., 2015). Together with our studies showing that Met-signaling is important for the migration of DC and thus for their immune function, it appears inevitable to address the immunological consequences of Met inhibition-based therapies in cancer patients.

## Acknowledgements

We are grateful to Ms. Katrin Götz for excellent technical assistance. We thank Dr. Corinna Rösseler for expert help in qPCR primer design. This work was supported by the Flow Cytometry Facility, a core facility of the Interdisciplinary Center for Clinical Research (IZKF) Aachen within the Faculty of Medicine at RWTH Aachen University. This work was funded in part by a grant from Deutsche Forschungsgemeinschaft (DFG HI1103/1-1 to TH).

## Notes

### Competing Interest Statement

The authors have declared no competing interest.

