## Supplemental Material for "Met-signaling Controls Dendritic Cell Migration by Regulating Podosome Formation and Function"

[1] Institute for Biomedical Engineering, Department of Cell Biology, RWTH Aachen University Medical School, Pauwelsstr. 30, 52074 Aachen, Germany
and
[2] Helmholtz-Institute for Biomedical Engineering, RWTH Aachen University, Pauwelsstr. 20, 52074 Aachen, Germany

### Supplemental Materials and Methods

*RNA isolation and quantitative reverse transcription - PCR (qPCR)*

RNA was isolated either using MagMAX-96 total RNA isolation kit (Life Technologies, USA) or NucleoSpin RNA kit (Macherey-Nagel, Germany) according to manufacturer's instructions. Isolated RNA was reverse transcribed using High Capacity cDNA Reverse Transcription Kit (Applied Biosystems, USA) according to the manufacturer's instructions. Then, 5-20 ng of cDNA were applied to quantitative PCR (qPCR) reaction plates with Fast SYBR Green Master Mix (Applied Biosystems, USA). The qPCR reaction was performed using a StepOnePlus Real Time PCR system (Applied Biosystems, USA). StepOne software v2.3 (Life Technologies, USA) was used to analyze the data by normalizing the cycle threshold (Ct) values of target genes to the corresponding Ct values of glyceraldehyde 3-phosphate dehydrogenase (*Gapdh*). Primer sequences are listed in supplemental Table 1 below.

**Suppl. Table 1. qRT-PCR primer sequences**

| Gene | Primer | Sequence (5´→ 3´) | Reference |
|---|---|---|---|
| *Gapdh* | forward<br>reverse | ACCTGCCAAGTATGATGACATCA<br>GGTCCTCAGTGTAGCCCAAGAT | (Tagoh et al., 2004) |
| *Cdh1* | forward<br>reverse | TGACGCAGCTCAAGAATCTC<br>TTGTTCTGGTTATCCGCGAG | This manuscript |
| *Epcam* | forward<br>reverse | GCGTGAGGACCTACTGGAT<br>CAAGCTCTGATGGTCGTAGG | This manuscript |
| *Mmp2* | forward<br>reverse | GACATACATCTTTGCAGGAGACAAG<br>TCTGCGATGAGCTTAGGGAAA | This manuscript |
| *Mmp9* | forward<br>reverse | CCTGGAACTCACACGACATCTTC<br>TGGAAACTCACACGCCAGAA | (Chen et al., 2005) |
| *Mmp14* | forward<br>reverse | CCCTAGGCCTGGAACATTCT<br>TTTGGGCTTATCTGGGACAG | This manuscript |
| *Ccr7* | forward<br>reverse | GTACGAGTCGGTGTGCTTC<br>GGTAGGTATCCGTCATGGTCTTG | (Moran et al., 2014) |
| *Cxcr4* | forward<br>reverse | AAACTGCTGGCTGAAAAGGC<br>TGACGTCGGCAAAGATGAAG | This manuscript |

### Supplemental Results

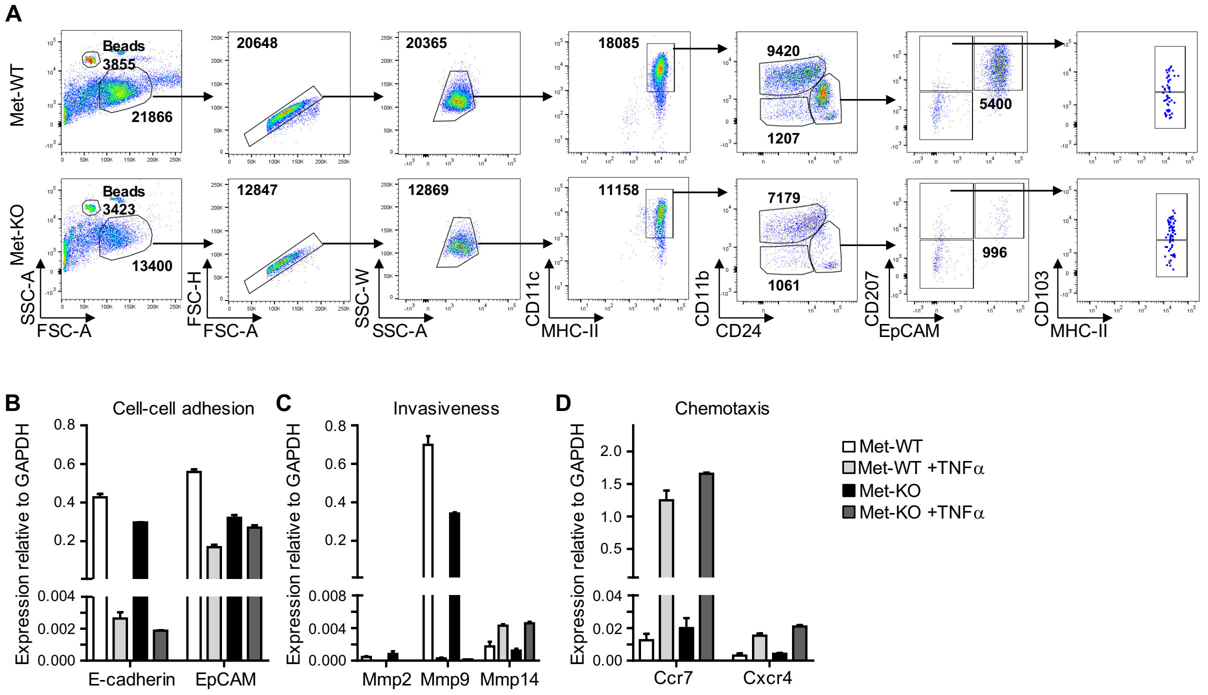


**Suppl. Fig. 1. Gating of emigrated Met-KO and Met-WT LC and dDC for flow cytometry analysis and gene expression analyses of sorted Met-KO and Met-WT LC.** (A) Ear dorsal halves from Met-KO and Met-WT mice were painted with acetone/DBP and cultured for 24 h and 48 h. Emigrated LC and dDC in medium were analyzed by flow cytometry. Gating strategy for dDC (CD11c+ MHC-II+), CD11b+dDC (CD11c+ CD11b+ CD24-), CD11b-dDC (CD11c+ CD11b- CD24-), CD103+dDC (CD11c+ CD11b- CD24+ CD207+ CD103+), CD103-dDC (CD11c+ CD11b- CD24+ CD207+ CD103-) and migratory LC (CD11c+ CD11bint CD24+ CD207+ EpCAM+). (B-E) Epidermal single cell suspensions from Met-KO and Met-WT skin were generated. LC were either FACS sorted immediately or after culturing for 24 h in the presence of TNFα. LC were pooled from two mice per experiment. Gene expression was quantified using qPCR done in triplicates. Bar graphs show the expression levels of genes associated with (B) cell-cell adhesion, (C) invasiveness, and (D) chemotaxis relative to the housekeeping gene *gapdh*. Error bars indicate SD.

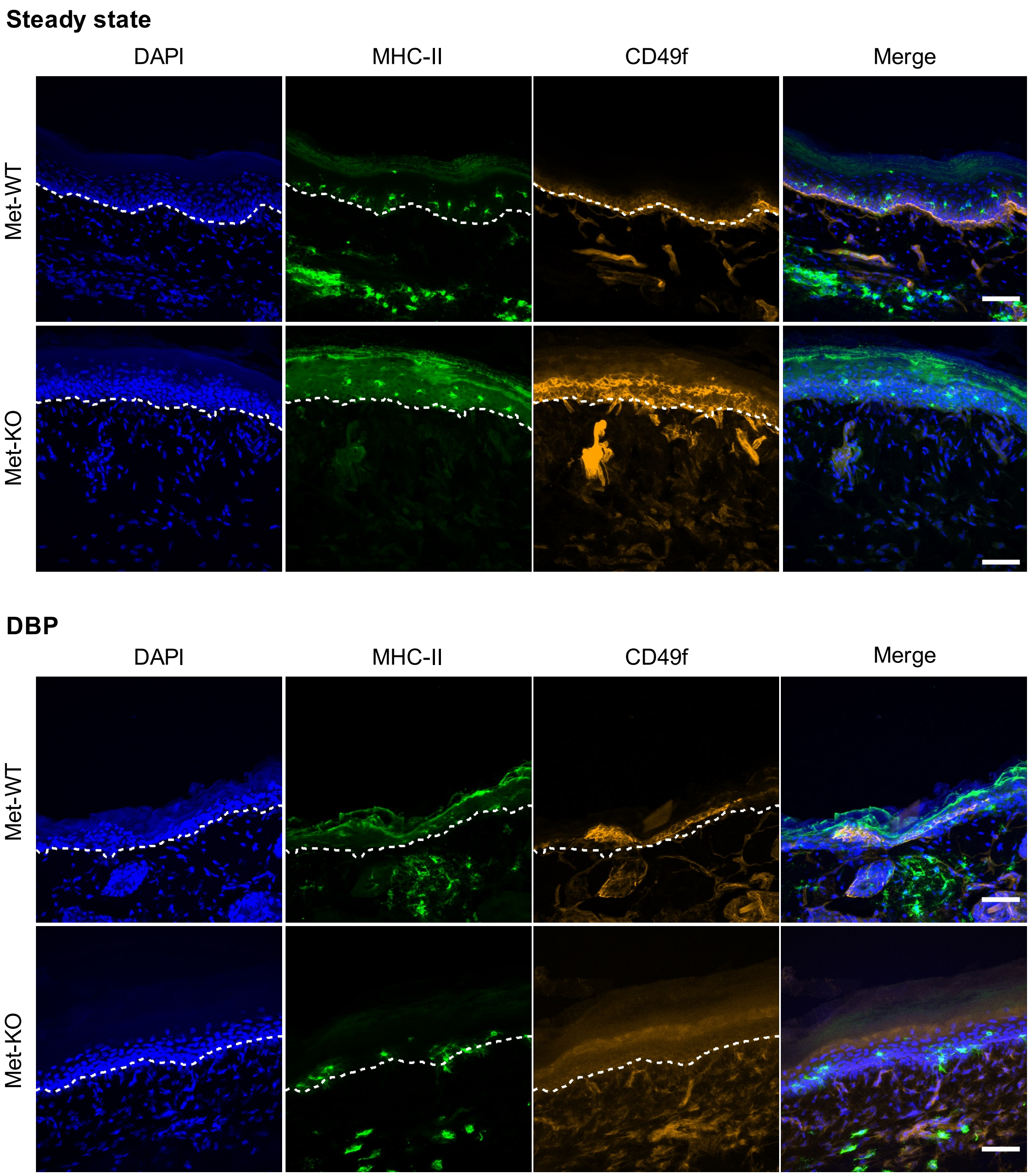


**Suppl. Fig. 2. Immunofluorescence analysis of Met-KO and Met-WT skin sections at steady state and after painting with acetone/DBP.** Footpad skin was isolated from Met-KO and Met-WT mice. Samples were either immediately cryopreserved in OCT or painted with acetone/DBP, cultured for 24 h and then cryopreserved in OCT. Then, the skin was sectioned and stained with anti-MHC-II, anti-CD49f and DAPI. Representative microscopic images of LC and dDC (green) in skin. Dotted line represents the epidermal dermal junction (EDJ). Scale bar, 50 μm.

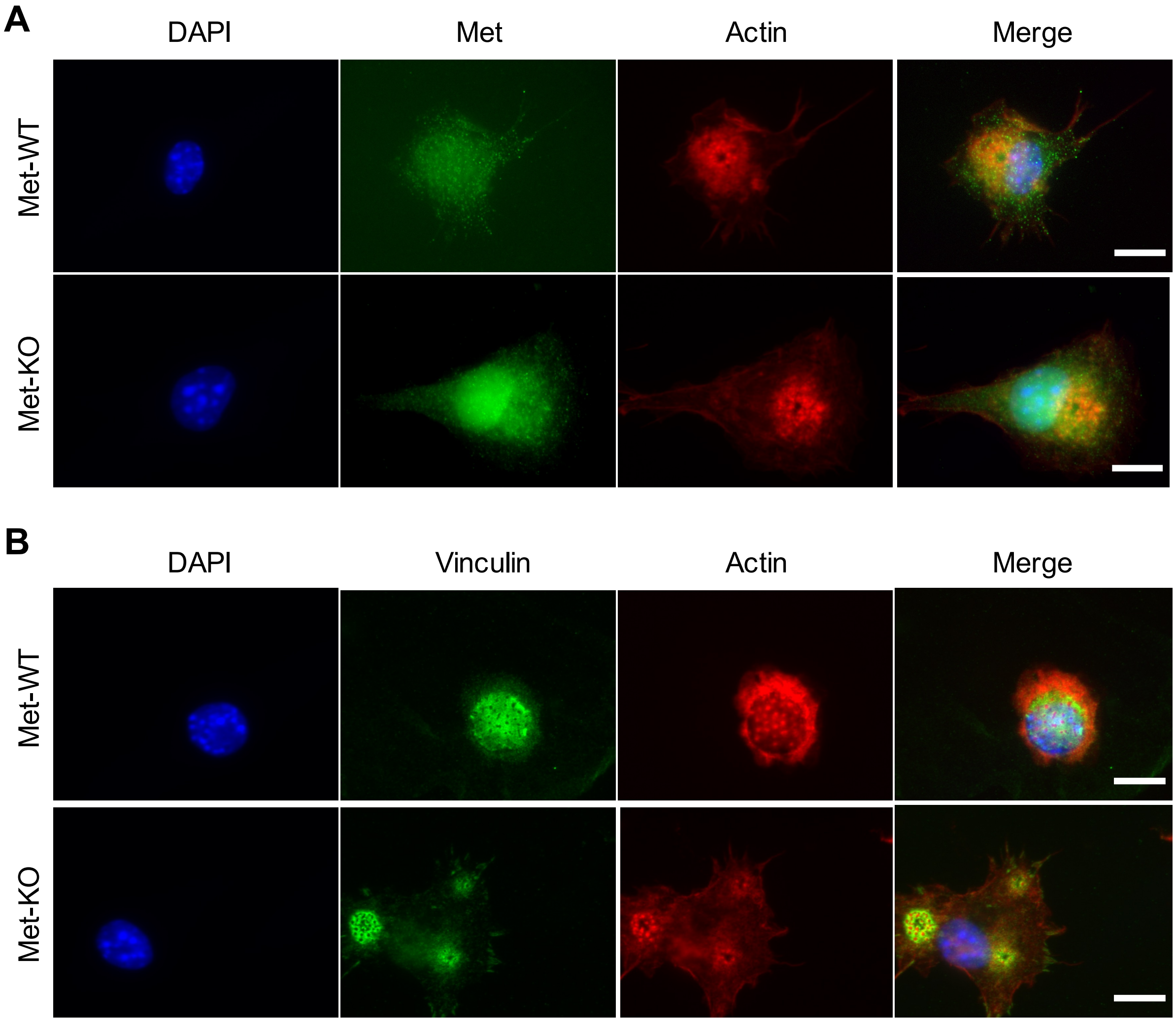


**Suppl. Fig. 3. Immunofluorescence analysis of podosomes in Met-KO and Met-WT BMDC.**
(A, B) Met-KO and Met-WT BMDC were incubated overnight on gelatin coated coverslips in the presence of TNFα. Then, BMDC were fixed and stained with DAPI (blue), phalloidin (red) and anti-Met (Cell Signaling, USA; clone C1D1) or anti-Vinculin (Sigma-Aldrich, USA; clone hVIN-1) Abs (green). Representative microscopic images of Met-KO and Met-WT BMDC BMDC forming podosomes with actin core cluster colocalizing with (A) Met and (B) Vinculin. Scale bar, 10 μm.

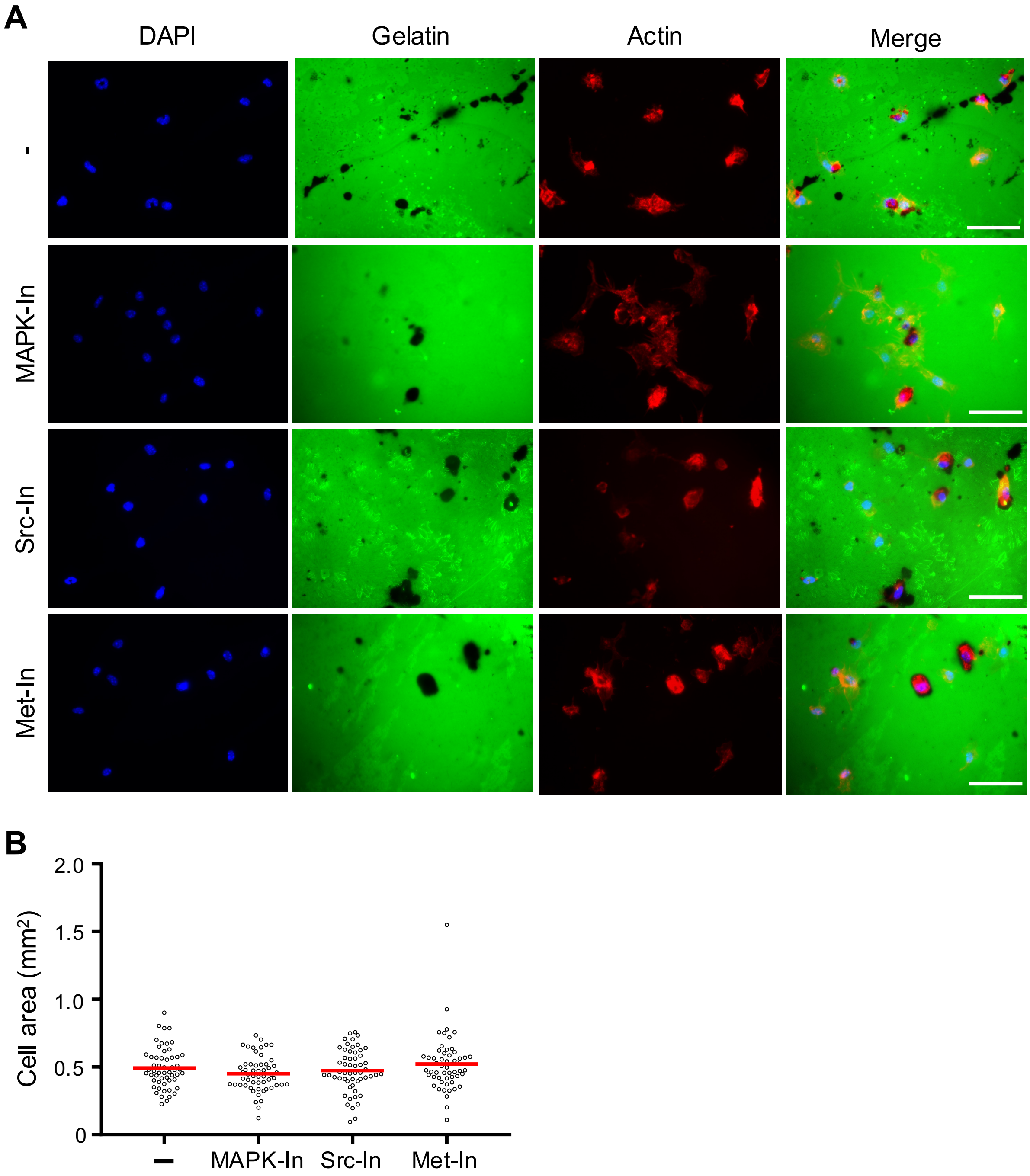


**Suppl. Fig. 4. Met-signaling impacts on podosome formation and ECM degradation in DCs.** Gelatin degradation and podosome formation assays were performed with Met-WT BMDC cultured on fluorescently labeled gelatin (green) for 24 h in the presence of TNF$\alpha$ and with or without 5 µM PD98059 (MAPK-In), 2 µM SU6656 (Src-In) or 1 µM SU11274 (Met-In). Then, BMDC were fixed and stained with DAPI (blue) and phalloidin (red). (A) Representative immunofluorescence microscopy images of gelatin degradation and podosome formation with untreated and treated BMDC. Scale bar, 50 µm. (B) Scatterplot shows cell area normalized to the number of cells per field. Data are from three independent experiments with at least 15 fields analyzed per experiment.

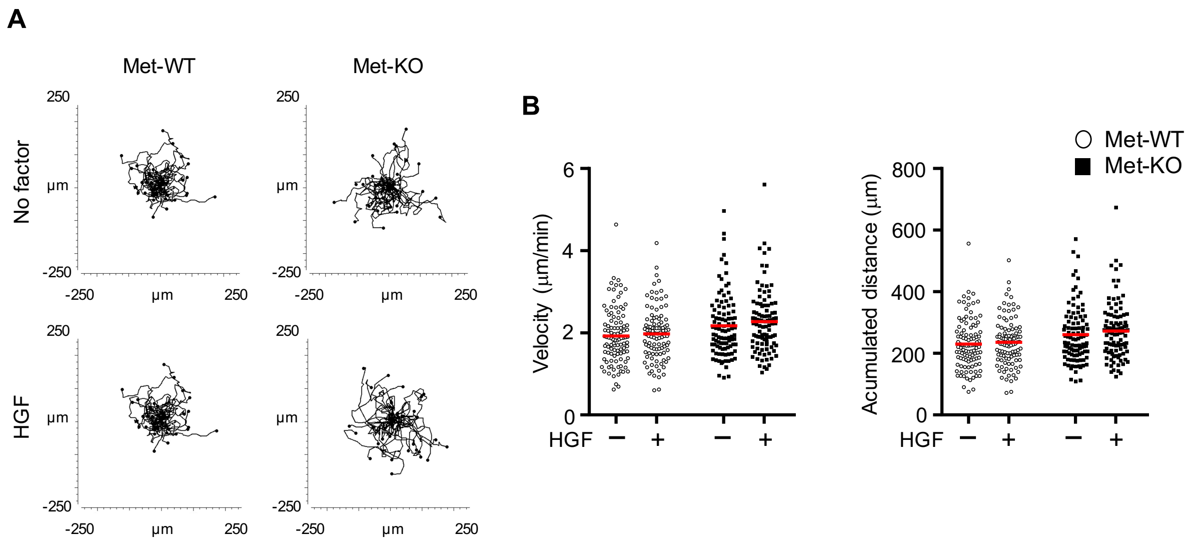


**Suppl. Fig. 5. Impact of Met-signaling on motility of immature DC in 3D collagen gel migration.** Immature Met-KO and Met-WT BMDC were either treated with HGF overnight or left untreated before they were embedded in collagen. Migration of embedded cells was visualized for 2 hours using time-lapse bright field microscopy. (A) Representative trajectory blots of Met-KO and Met-WT BMDC in the absence or presence of HGF. Each dot represents one cell. (B) Scatter plots show velocity (left panel) and accumulated distance from starting point (right panel). Data were pooled from three independent experiments with ≥ 30 cells for each condition per experiment.

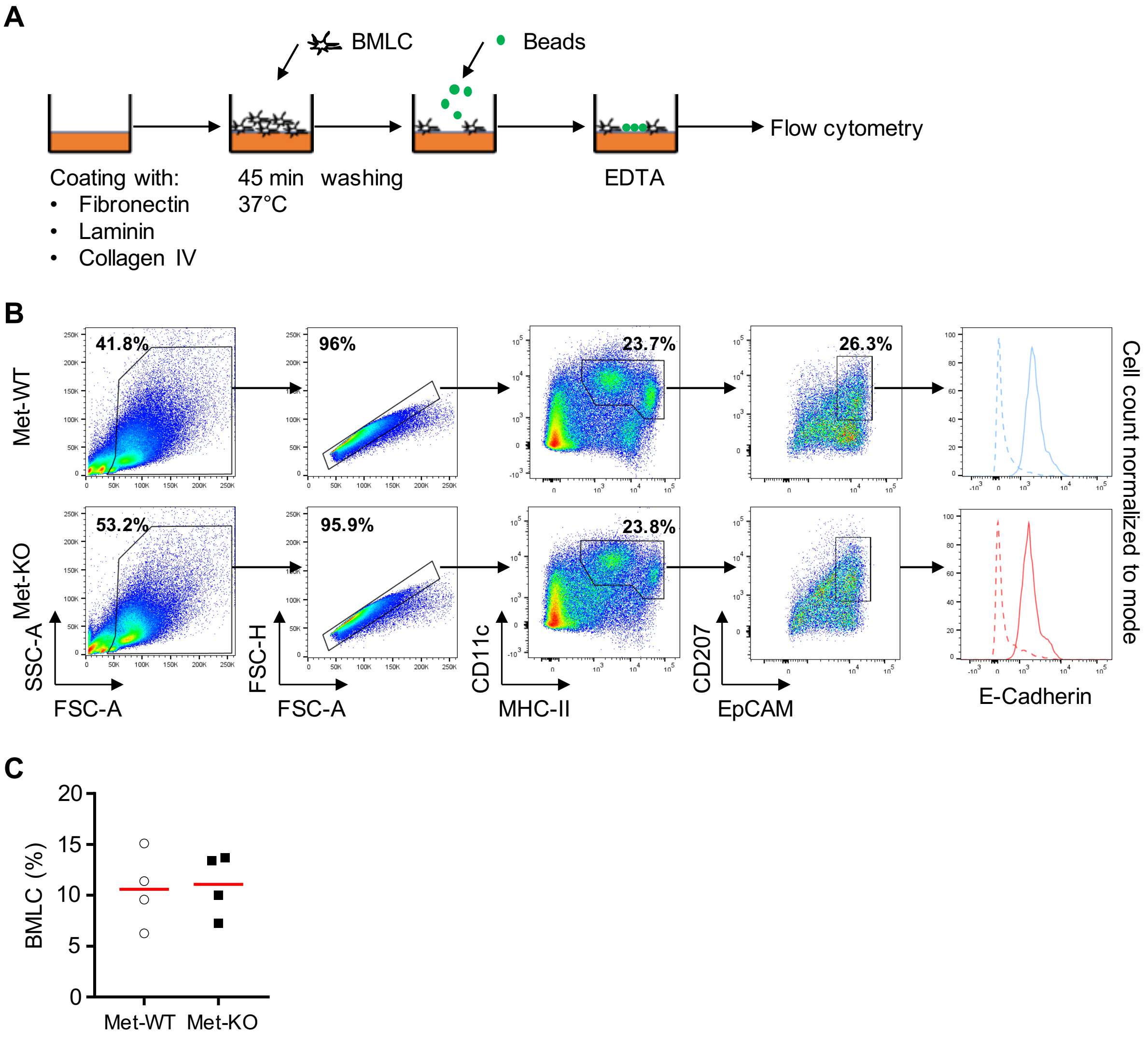


**Suppl. Fig. 6. Adhesion of Met-KO and Met-WT BMLC to ECM components.** TGF-β-derived BMLC were generated from Met-KO and Met-WT BM and treated with or without HGF overnight. Then, BMLC adhesion to ECM factors fibronectin, laminin and collagen IV was assessed. (A) Schematic representation of the experimental procedure. (B) Representative flow cytometry analysis of TGF-β-derived BMLC (CD11c+ MHC-II+ CD207+ EpCAM+) expressing E-cadherin generated from Met-KO and Met-WT BM. Solid lines, stained; dashed lines, unstained. (C) Scatter plot shows the percentage of Met-KO and Met-WT BMLC in cell culture at day 5 of differentiation. Data shown are from four independent experiments.
